# Stiffness anisotropy coordinates supracellular contractility driving long-range myotube-ECM alignment

**DOI:** 10.1101/2023.08.08.552197

**Authors:** Nathaniel P. Skillin, Bruce E. Kirkpatrick, Katie M. Herbert, Benjamin R. Nelson, Grace K. Hach, Kemal Arda Günay, Ryan M. Khan, Frank W. DelRio, Timothy J. White, Kristi S. Anseth

## Abstract

In skeletal muscle tissue, injury-related changes in stiffness activate muscle stem cells through mechanosensitive signaling pathways. Functional muscle tissue regeneration also requires the effective coordination of myoblast proliferation, migration, polarization, differentiation, and fusion across multiple length scales. Here, we demonstrate that substrate stiffness anisotropy coordinates contractility-driven collective cellular dynamics resulting in C2C12 myotube alignment over millimeter-scale distances. When cultured on mechanically *anisotropic* liquid crystalline polymer networks (LCNs) lacking topographic features that could confer contact guidance, C2C12 myoblasts collectively polarize in the stiffest direction of the substrate. Cellular coordination is amplified through reciprocal cell-ECM dynamics that emerge during fusion, driving global myotube-ECM ordering. Conversely, myotube alignment was restricted to small local domains with no directional preference on mechanically *isotropic* LCNs of same chemical formulation. These findings reveal a role for stiffness anisotropy in coordinating emergent collective cellular dynamics, with implications for understanding skeletal muscle tissue development and regeneration.

## Introduction

Mechanotransduction and cell migration play critical roles in developmental, homeostatic, regenerative, and pathological biological processes.[1,2] Early *in vitro* studies identified preferential cellular migration up a stiffness gradient using polyacrylamide hydrogels, a process termed durotaxis.[3] Recently, Sunyer *et al.* showed that while MCF10A cells did not durotax in isolation on polyacrylamide gels with a stiffness gradient >50 kPa/mm, clusters of cells migrated together towards stiffer gel regions in a form of collective cell durotaxis.[4] The enhanced sensitivity of MCF10A monolayers to mechanical cues was found to be dependent on long-range force transmission through cell-cell junctions as well as cellular contractility, which effectively allowed cells to sense a stiffness gradient as a collective unit. Collective durotaxis has also been observed *in vivo*, where *Xenopus laevis* neural crest cells were found to follow a receding stiffness gradient during development, which enforced supracellular polarity of cell-matrix adhesions throughout the migrating cell cluster.[5]

While cellular responses to stiffness *gradients* are relatively well understood, there is also strong precedent that stiffness *anisotropy* can serve as a mechanical cue influencing cellular behaviors such as polarization. Bischofs and Schwarz introduced a connected theoretical framework to explain the observed polarization of single cells on anisotropic substrates, whereby focal adhesions (FAs) aligned in the stiffest direction mature more rapidly and outcompete FAs in other directions due to energetic favorability, causing cells to polarize with the stiffest direction of the substrate.[6] This theory has been validated experimentally with isolated cells polarizing in the stiffest direction on strained collagen hydrogels as well as soft (∼2 MPa) anisotropic polydimethylsiloxane micropillars.[7,8] However, when the same cells were grown on anisotropic polystyrene micropillars with a much higher baseline elastic modulus (∼2 GPa), no preferential cell polarization was observed.[7] Thus, if a mechanically anisotropic substrate has sufficiently supraphysiological stiffness in all directions, the relative energetic favorability of FA maturation in the stiffest direction is diminished and isolated cells will polarize randomly.[6] While single cell responses to stiffness anisotropy may be limited by this mechanism, the boundaries of collective cellular responses to stiffness anisotropy have yet to be explored. This gap in understanding exists in part because there are limited strategies for generating mechanically anisotropic substrates that lack anisotropic topography (e.g., fibrous materials), which introduces confounding effects of contact guidance.

Liquid crystalline polymer networks (LCNs) are emerging as a novel class of anisotropic biomaterials for tissue engineering, medical devices, and soft robotics. Typical nematic liquid crystalline molecules (mesogens) contain a rigid rod-like aromatic core flanked by one or more reactive end groups.[9] Molecular alignment of reactive mesogens can be enforced by surface, shear, and optical patterning methods due to strong intermolecular interactions and is retained upon crosslinking the mesogens into a network.[10–12] As a result of the flexibility of approaches developed to program molecular anisotropy into a polymer network, there is growing interest in engineering LCNs to control tissue morphology. For example, Turiv *et al.* used photoalignment techniques to synthesize LCNs with spatial control of ellipsoidal surface features, which guided fibroblast polarization to closely match the user-defined photopattern.[13] Still, *in vitro* strategies to program cellular alignment that rely on topographic features (e.g., grooves) for contact guidance do not reflect the true nature of mechanobiological cues that cells harness to (re)generate aligned tissues *in vivo*.

In cases where mesogen alignment is programmed to be unidirectional across a large area, polymerization of reactive end groups results in a monodomain LCN (mLCN) that is stiffest in the direction of molecular alignment, also known as the nematic director.[14] Recently, fibroblasts cultured on mLCNs lacking topographic features were found to align with the nematic director.[15] Notably, Martella *et al.* also observed that C2C12 myoblasts cultured on mLCNs lacking topographic features aligned with the nematic director.[16] However, the cellular dynamics involved in myoblast alignment were not addressed, and none of these studies have systematically explored whether changing the degree of substrate anisotropy impacted alignment. LCN moduli and stiffness anisotropy can be tuned by altering the ratio of crosslinker to chain extender,[10] thereby offering a flexible platform to probe the mechanoregulation of collective cellular responses to stiffness anisotropy *in vitro*.

Much like mLCNs, muscle tissue is inherently anisotropic with greater elastic modulus in the longitudinal direction compared to the transverse direction.[17,18] It is well known that the highly anisotropic mechanics and architecture of the extracellular matrix (ECM) play critical roles in myoblast fusion and alignment during development and regeneration.[19] In zebrafish craniofacial development, myocytes initially attach to cartilage before bone growth starts.[20] Cartilage then undergoes directional expansion, which applies stretch-induced tension on myocytes that drives polarization in the correct orientation before fusion begins. When laminin subunits were knocked out, myocytes were unable to polarize correctly, and myofibrils were found to be misaligned. Upon injury, muscle stem cells (MuSCs) sense changes in ECM stiffness and initiate mechanosensitive signaling pathways that activate and maintain a population of proliferating MuSCs.[21,22] MuSCs then rely on the anisotropic basal lamina as a template to effectively repair small defects with functional, aligned muscle fibers,[23] as repair of defects lacking an intact basal lamina results in randomly oriented myofibers that cannot effectively generate force.[19] Intravital imaging of skeletal muscle regeneration in mice demonstrated that myoblasts polarize with the longitudinal fiber direction inside these basal lamina remnants before fusing into myotubes.[24] However, the specific biophysical cues that direct myoblast polarization prior to fusion, as well as the collective cellular dynamics responsible for organizing aligned skeletal muscle tissue over large distances, remain unclear. We hypothesized that stiffness anisotropy acts as a mechanobiological cue for collective myoblast polarization and myotube alignment. To address this question, we cultured C2C12 murine myoblast cells on mLCNs and interrogated the collective cellular dynamics that emerged over increasing length scales.

Here, we demonstrate that C2C12 myoblasts cultured on mLCN substrates lacking topography adhere and polarize randomly until reaching a critical cell density threshold, after which supracellular coordination of actomyosin contractility develops resulting in global myotube and ECM alignment with the nematic director over large distances (∼ millimeters). We found that myotube alignment was enhanced on stiffer mLCNs with greater orthogonal difference and further scaled with the degree of stiffness anisotropy. Live-cell imaging revealed the combination of crowding dynamics, collective myoblast polarization, and contractile cellular flows mediated by reciprocal cell-ECM dynamics synergize to yield long-range alignment of myotubes with the nematic director. This study of cellular responses to stiffness anisotropy across multiple length scales using novel anisotropic substrates adds to the emerging importance of collective cellular dynamics as a complement to molecular and genetic mechanisms in tissue development and regeneration.

## Results

### LCN films prepared by aza-Michael addition reaction exhibit distinct mechanical properties

LCNs are commonly prepared via photopolymerization of liquid crystalline monomers. Recent reports detail the utilization of chain-extending oligomerization reactions to adjust the glass transition temperature and associated moduli of LCNs.[10,11] Motivated by the potential to study the collective cellular response of myoblasts to stiffness anisotropy, we focus on monodomain LCNs prepared by surface-mediated alignment. Accordingly, we prepared mLCNs via an aza-Michael addition reaction at acrylate molar fraction of 0.5, 0.75, and 1.0 to fabricate cell culture substrates with orientational order over a range of moduli **(Figures 1A and S1A–C; Table S1)**. Isotropic LCNs (iLCNs) with the same chemical formulations but lacking stiffness anisotropy were prepared as controls. mLCN alignment was confirmed with polarized optical microscopy **(Figure S1D)**. Tensile testing parallel and orthogonal to the nematic director was used to measure the magnitude of the elastic moduli as well as the degree of stiffness anisotropy between the three mLCN formulations **(Figures 1B and S1E)**. Isotropic LCNs exhibit isotropic moduli in between the parallel and perpendicular elastic modulus for monodomain samples of the same formulation.[25] As expected, the mLCN elastic moduli parallel to the nematic director trended with crosslink density (mLCN-0.5 < 0.75 ∼ 1.0).[10] Meanwhile, the stiffness anisotropy ratio (E_||_ / E_1−_) was highest for mLCN-0.5 (4.5x), while the orthogonal difference in elastic modulus (E_||_ – E_1−_) was highest for mLCN-1.0 (900 MPa). mLCN-0.75 provided a unique comparison to mLCN-1.0 as the elastic moduli parallel to the nematic director were almost identical (∼1.4 GPa), while the degree of stiffness anisotropy was greater for mLCN-1.0 **(Figure 1B)**.

**Figure 1.**
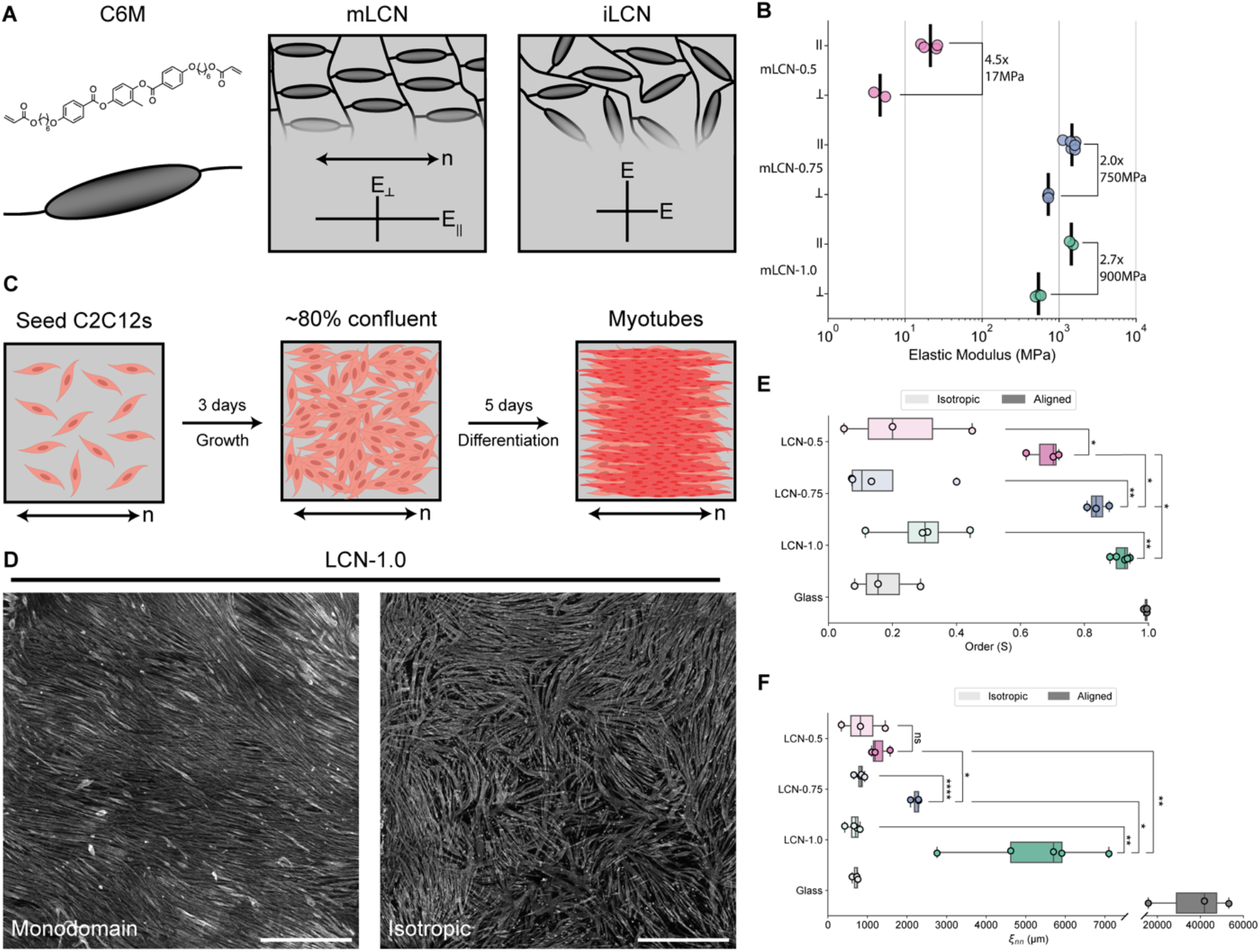
mLCN stiffness anisotropy drives C2C12 myotube ordering. (A) Illustration of monodomain and isotropic LCN network structure and mechanical anisotropy (n indicates nematic director of mLCN). **(B)** Elastic modulus parallel and orthogonal to mLCN nematic director derived from the initial linear regime of stress-strain curves. Stiffness anisotropy ratio (E_||_ / E_λ_) and difference (E_||_ – E_λ_) is calculated from the mean of repeated tensile tests. Line indicates mean, all replicates shown. **(C)** Illustration of C2C12 growth and differentiation on mLCNs. Created with BioRender.com. **(D)** Representative images of myotubes stained for myosin II heavy chain (MF-20, gray) after 5 days of differentiation on monodomain (left) and isotropic (right) LCN-1.0. Scale bars = 1000 µm. **(E)** Myotube orientation-order parameter (S) and **(F)** nematic correlation length (µm) of myotubes after 5 days of differentiation on isotropic and aligned substrates (isotropic vs. monodomain for LCNs, glass coverslip vs. NanoSurface substrate for Glass). For the box plots in **(E)** and **(F)**, the box limits extend from the 25^th^ to 75^th^ percentiles; the horizontal line indicates the median value; the whiskers extend by 1.5x the inter-quartile range; all replicates including outliers are shown. Statistical analysis was performed with one-tailed (isotropic v monodomain) or two-tailed (monodomain v monodomain) unpaired Student’s *t*-test with Welch’s correction with significance claimed at *p < 0.05, **p < 0.01, ***p < 0.001. ****p < 0.0001. See also **Figures S1, S2, S3, and S4; Table S1**.

We then characterized the surface of mLCNs with atomic force microscopy (AFM) to assess for the presence of any topographic features that might confer contact guidance **(Figures S1G and S1H)**. All compositions had similar root mean square (rms) roughness measurements of ∼2 nm. However, height maps of mLCN-0.75 and mLCN-1.0 indicated the presence of some nanoscale raised defects with a maximum height of ∼10 nm, which were not anisotropic in their structure or distribution **(Figure S1G)**. Many prior studies document the importance of the depth of topographical features in aligning cells, with a threshold of ∼35 nm required for contact guidance.[26,27] Therefore, the nanoscale features on these mLCNs were well below this threshold for imparting topographic guidance and are not expected to contribute to cell alignment. Finally, contact angle measurements showed little difference in surface energies between the three compositions with all materials being slightly hydrophilic with contact angles ranging from 74– 83° **(Figure S1F)**.

### Myotube alignment is influenced by the degree of mLCN stiffness anisotropy

Following mLCN film preparation, we seeded C2C12 myoblasts at a relatively low density (5,000 cells/cm^2^) on sterilized mLCNs as well as gelatin-coated glass coverslips for an additional isotropic, stiff control used in myoblast culture.[28] Consistent with previous reports, we observed that C2C12s adhered to neat mLCNs, presumably due to non-specific protein adsorption from the growth media.[13,16,29] Initially, we assessed whether isolated myoblasts would preferentially polarize with the nematic director. When myoblast orientation was quantified across the first three days of growth, there were no differences between mLCNs and glass coverslips **(Figures S2A and S2B)**. Furthermore, there were minimal differences among other measured cellular outputs (i.e., proliferation rate, cell area, aspect ratio) before confluence **(Figures S2C–E)**. We attribute these similarities to the high overall stiffness (MPa–GPa) of all three mLCN compositions,[6,7] and establish that both isolated myoblasts and small clusters of myoblasts cannot sense the stiffness anisotropy of these mLCNs.

After three days in growth media, C2C12 myoblasts were ∼80% confluent, at which point myoblast differentiation and fusion was induced by serum starvation **(Figure 1C)**.[30] Samples were fixed and immunostained after three and five days of differentiation, representing the stages of nascent myotube formation and myotube maturation, respectively.[31,32] Prolonged culture times exceeding 6 days after the induction of differentiation were difficult to maintain, as myotubes became highly contractile and detached from their substrates, which has been previously observed in similar cultures.[30] Myotube alignment (with a reference of 0° matching the orientation of the nematic director of the mLCN), 2D orientation-order parameter (S, based on the 3D liquid crystalline nematic order parameter), and nematic correlation length (ξ_nn_), were quantified using OrientationJ and custom scripts (see STAR Methods).[28,33]

After three days of differentiation, evidence of myotube alignment with the nematic director was already present on all three mLCN formulations **(Figure S3)**. After five days of differentiation, myotubes cultured on mLCNs were highly aligned, whereas myotubes differentiated on iLCNs and glass displayed some regions of local alignment, but no long-range alignment **(Figures 1D and S3A)**. Unexpectedly, the mean orientation of myotube alignment on mLCNs was consistently offset ∼10–20° clockwise (CW) from the nematic director **(Figures S3A and S4; Supplementary Note 1)**. Myotube order and ξ_nn_ increased between day three and day five on all mLCNs, while only ξ_nn_ increased on iLCNs and glass due to continued fusion and elongation of myotubes **(Figures S3B and S3C)**.[28] Notably, both myotube order and ξ_nn_ were significantly higher on all mLCNs compared to their formulation-matched iLCN controls, with the exception of ξ_nn_ for mLCN-0.5 **(Figure 1E)**. This direct comparison between C2C12 myotubes cultured on monodomain and isotropic LCNs is strong evidence that C2C12s collectively sense the stiffness anisotropy of mLCNs during differentiation and align with the stiffest direction. Furthermore, myotube order after five days of differentiation was significantly higher on the stiffer mLCNs (S = 0.84 (mLCN-0.75), 0.92 (mLCN-1.0)) compared to the much softer mLCN-0.5 (S = 0.68) **(Figure 1E)**. Myotube order was also significantly greater on mLCN-1.0 compared to mLCN-0.75. Impressively, myotube order on mLCN-1.0 approached the order of myotubes grown on a commercial polymeric substrate with linear grooves used as a positive control for myotube alignment (NanoSurface, S = 0.99). The length scale of myotube alignment on mLCNs followed a similar trend with mean ξ_nn_ values of 1300, 2200, and 5200 µm for mLCN-0.5, mLCN-0.75, and mLCN-1.0, respectively **(Figure 1F)**.

### Stiffness anisotropy coordinates collective cellular dynamics across multiple length scales

To elucidate the spatiotemporal dynamics of myotube alignment on mLCNs, we employed live-cell imaging with fluorescent DNA and actin probes and assessed the evolution of myoblast and myotube ordering. We primarily focused on LCN-1.0 as the anisotropic properties of mLCN-1.0 had the greatest effect on myotube alignment. Hereafter, these substrates are simply referred to as mLCNs and iLCNs. While we already established that single cells and clusters of cells did not align with the nematic director on mLCNs, we hypothesized that alignment could be induced by preferential migration of myoblasts in line with the nematic director above a critical cell density threshold. Although we characterized only the parallel and orthogonal mLCN modulus values, the mechanical properties are better described as a radial stiffness gradient;[34] analogous to the linear stiffness gradient of the polyacrylamide gels used by Sunyer *et al.* that induced collective durotaxis.[4,35] Myoblast migration was evaluated at the single-cell level with nuclear tracking until the cell density became too high for accurate segmentation (>2200 cells/mm^2^). As expected, migration speed continuously decreased during the proliferative phase, as mobility became increasingly limited due to cellular crowding **(Figure 2B)**.[36] However, we found no strong evidence of orientationally-biased C2C12 division **(Figure S5)** or preferential migration **(Figure 2B)**.

**Figure 2.**
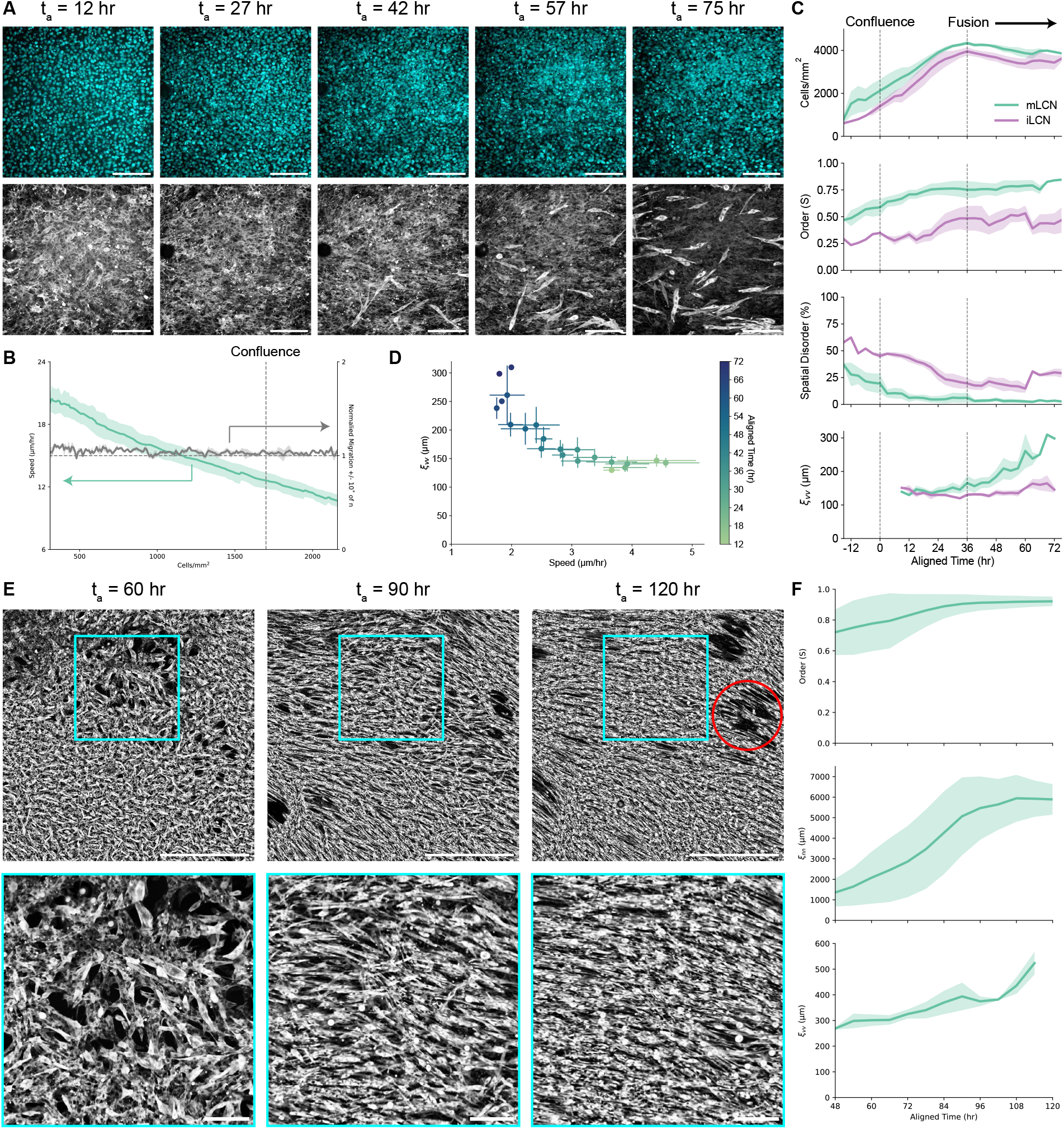
Stiffness anisotropy coordinates collective cellular dynamics across multiple length scales. (A) Representative images of C2C12 nuclei (top) and actin (bottom) on mLCNs at t_a_ = 12, 27, 42, 57, and 75 hr (left to right). Scale bars = 200 µm. **(B)** Myoblast migration speed (µm/hr; left axis) and normalized migration ± 10° from nematic director (right axis) on mLCNs. Dashed line indicates approximate density at which myoblasts achieved confluence. Data are represented as mean ± SEM. **(C)** Top to bottom: Temporal evolution of cell density (cells/mm^2^), orientation-order parameter (S), spatial disorder (%), and velocity correlation length (µm) on mLCNs and iLCNs. Dashed lines indicate approximate time at which myoblasts achieved confluence (left) and initiated fusion (right). Data are represented as mean ± SEM. **(D)** Velocity correlation length (µm) as a function of cellular speed (µm/hr) after confluence on mLCNs. Data are represented as mean ± SEM. **(E)** Representative images (top) and inset (bottom) of C2C12 actin on mLCNs at t_a_ = 60, 90, and 120 hr (left to right). Scale bars = 1000 µm (top) and 200 µm (bottom). Red circle indicates location of a contractile defect. **(F)** Top to bottom: Temporal evolution of orientation-order parameter (S), nematic correlation length (µm), and velocity correlation length (µm) on mLCNs. Data are represented as mean ± SEM. See also **Figures S5; Videos S1 and S2.**

We then performed another set of live-cell imaging experiments on both mLCNs and iLCNs starting at ∼80% confluence and ending after fusion of nascent myotubes **(Figure 2A; Video S1)**. Due to small experimental variation in initial seeding density, individual timeseries were aligned to each other in time (aligned time, t_a_) based on the timepoint at which myoblasts became confluent (t_a_ ∼ 0 hr; see STAR Methods). Myoblasts continued proliferating for an additional 36 hours past confluence, forming a multilayered culture with a maximum cell density of ∼4000 cells/mm^2^ on both mLCNs and iLCNs **(Figure 2C)**.[37] Myoblast order on both substrates was low (S ≤ 0.5) at the start of imaging, complementing our previous analysis that found no preferential polarization of myoblasts with the nematic director after three days of growth **(Figures 2C and S2B)**. However, we found that myoblasts collectively polarized with the stiffest direction of mLCNs as they approached confluence and continued proliferating **(Figure 2A; Video S1)**. Once myoblasts reached confluence at t_a_ = 0 hr, order on mLCNs had increased to 0.59 and continued to rise to 0.75 through t_a_ = 36 hr as myoblasts collectively polarized with the nematic director **(Figure 2C)**. Meanwhile, order remained below 0.5 on iLCNs through t_a_ = 36 hr. Spatial disorder, which was calculated as the percent of local regions (∼50 x 50 µm) with an order parameter less than 0.5, decreased from 37% to just 6% on mLCNs from t_a_ = -15–36 hr. In contrast, spatial disorder remained above 20% at t_a_ = 36 hr on iLCNs **(Figure 2C)**. Therefore, we posit that myoblasts are able to collectively sense the stiffness anisotropy of the mLCN above a critical cell density threshold that allows them to coordinate cytoskeletal contractility and gradually polarize with the stiffest direction of the mLCN.

Next, we investigated whether this collective polarization was caused by collective C2C12 migration after confluence using particle image velocimetry (PIV) analysis of the actin signal **(Video S1)**. Akin to the nematic correlation length (ξ_nn_), we calculated the velocity correlation length (ξ_vv_) from the angle of velocity vectors as a measure of the average length scale of collective migration. ξ_vv_ remained below ∼150 µm on both substrates from t_a_ = 9 hr until the end of the proliferative phase at t_a_ = 36 hr, confirming that the collective myoblast polarization seen on mLCNs was not caused by preferential cellular migration **(Figure 2C)**. Proliferation was then arrested as myoblasts began to fuse, forming nascent multinucleated myotubes. Over the next 39 hours (t_a_ = 36–75 hr), a collective cellular flow polarized with the nematic director was evident on mLCNs, corresponding to an increase in ξ_vv_ from ∼150 µm to ∼300 µm **(Figure 2C; Video S1)**. C2C12 order increased from 0.75 to 0.85 and spatial disorder fell to just 2.8% on mLCNs from t_a_ = 36–75 hr as cellular alignment was further enhanced by collective cellular migration. In contrast to epithelial monolayers that undergo jamming transitions resulting in glassy dynamics with low velocity correlation lengths,[36] C2C12 velocity correlation length increased with decreasing speed over time **(Figure 2D)**. While cellular flows also emerged on iLCNs during fusion, they were not polarized in any single direction and thus ξ_vv_ did not significantly increase. C2C12 order on iLCNs fluctuated around 0.5 during these randomly oriented cellular flows on iLCNs, yet spatial disorder increased from 20% to 30% illustrating that randomly oriented cellular flows impede the development of global cellular alignment **(Figure 2C)**. In sum, collective myoblast polarization in the stiffest direction of mLCNs precedes emergent collective cellular flows, which serve to further homogenize cellular alignment after collective polarization.

Examining the time series more closely, most of the nascent myotubes arising from myoblast fusion are already polarized with the nematic director on mLCNs **(Figure 2A; Video S1)**. This might be expected as myoblasts were already polarized before fusion began and have a well-documented preference for anisotropic fusion.[31] However, we also observed nascent myotubes that were not initially polarized with the nematic director became increasingly aligned by the polarized flow field. To further elucidate the size scale of these collective cellular flows, we performed a similar experiment but imaged live cells with the fluorescent actin probe across the entire ∼3x3 mm mLCN from t_a_ = 48–120 hr **(Figure 2E; Video S2)**. Using the same sequence of experiments and analysis routines, we again found that myoblast order was already high (S ∼ 0.7) before the cellular flows began **(Figure 2F)**. As nascent myotubes arose, local 2D orientation- order parameter analysis identified the presence of a few disordered regions of myotubes highlighted by the presence of topological defects that can be visualized using line integral convolution **(Video S2)**. As the length scale of the collective cellular flows increased from ξ_vv_ = ∼270 µm to ∼520 µm **(Figure 2F)**, these topological defects annihilated and disordered myotubes were reoriented by the flow field to generate a highly ordered myotube culture (S = 0.92) that matched our results for myotube order in **Figure 1E (Figure 2F; Video S2)**. Meanwhile, ξ_nn_ increased from 1800–6000 µm, illustrating the dramatic effect of the collective cellular flows inducing global myotube alignment **(Figure 2F)**. Taken together, these results indicate that in the absence of stiffness anisotropy, differentiating myoblasts cannot coordinate myoblast polarization nor the collective cellular flows that arise during fusion, leading to polydomain myotube alignment. In the presence of stiffness anisotropy, even when it is indistinguishable to single cells, myoblasts collectively polarize in the direction of greater stiffness. Subsequent collective cellular flows remain polarized with the nematic director and drive global myotube alignment on mLCNs.

### Reciprocal cell-ECM dynamics mediate collective cellular flows

To this point, we have established that collective myoblast polarization clearly plays a role in templating the collective cellular flows prior to the onset of differentiation on mLCNs. However, our observations and previous *in vitro* studies have demonstrated that fused myotubes lie on top of a layer of myoblasts.[38] Thus, nascent myotubes may not sense the stiffness anisotropy of the substrate, alluding to the contribution of an additional collective cellular dynamic. Given the importance of the ECM in muscle development and regeneration, we assessed whether dynamic cell-ECM interactions were mediating the increasing scale of these collective cellular flows. We first fixed and stained myoblasts grown on mLCNs shortly after they achieved confluence and found that a substantial amount of nascent fibronectin had already been deposited **(Figure 3A)**. Fibronectin alignment was quantified using OrientationJ, which revealed modest alignment with the nematic director that mirrored the myoblast actin alignment **(Figure 3A)**. This finding suggests that myoblasts are actively remodeling the ECM during the process of collective polarization. Subsequently, we examined the architecture of three key ECM proteins present in muscle tissue[39]— fibronectin, laminin, and collagen IV — after five days of differentiation on mLCNs. The alignment of all three cell-secreted ECM proteins was dramatically higher compared to the fibronectin staining before fusion and closely matched the corresponding myotube alignment in both magnitude and direction **(Figures 3B–D)**. These results clearly demonstrate that both cell and ECM alignment evolve together over time, but whether these cell-ECM dynamics mediate the polarized cellular flows was still unclear.

**Figure 3.**
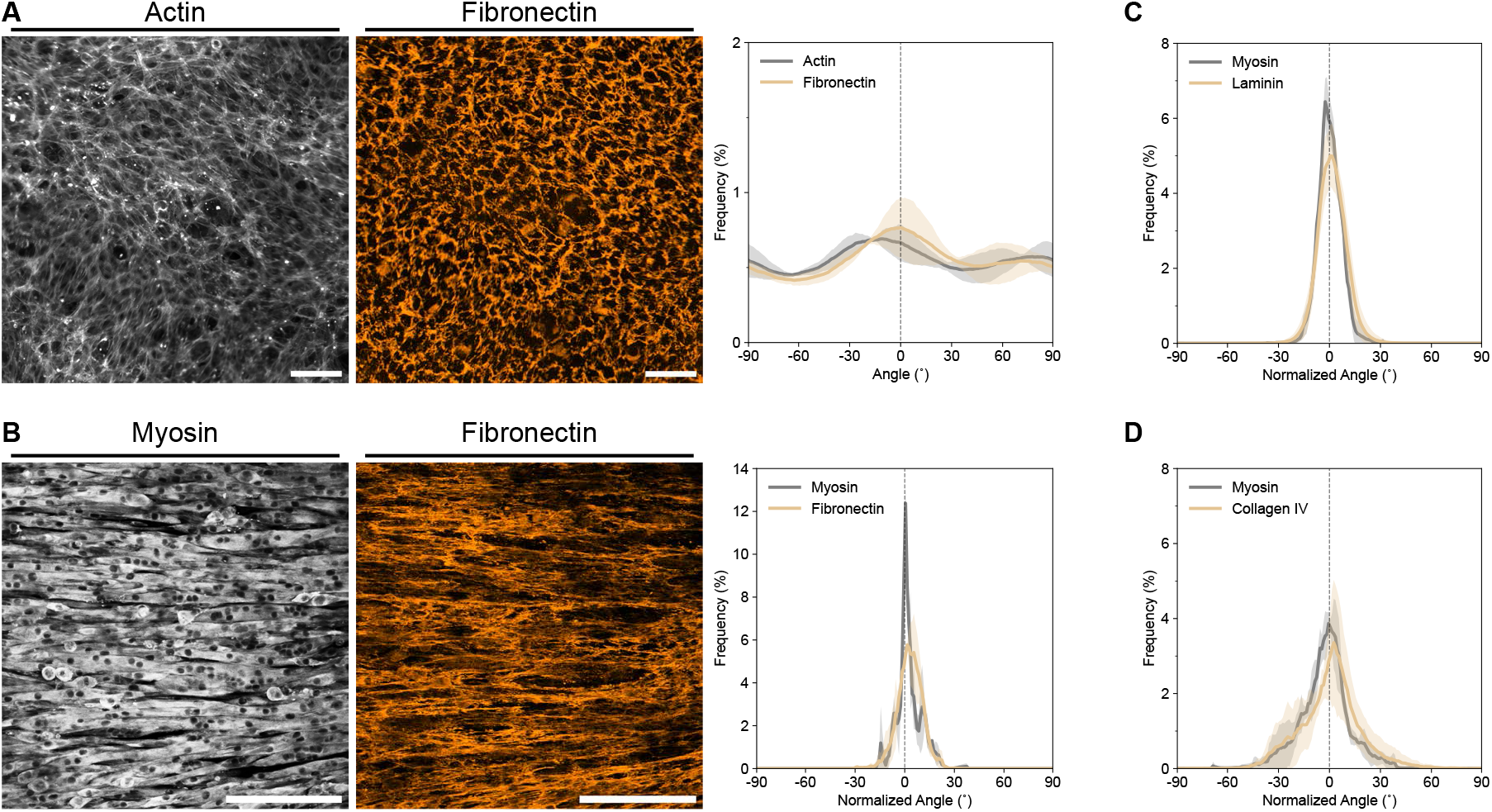
Nascent ECM alignment develops in parallel with myotube alignment. (A) Representative maximum intensity projections of actin (left) and fibronectin (middle) on mLCNs after reaching confluence. Scale bars = 100 µm. Corresponding frequency distribution plot (right) of actin and fibronectin alignment. Data are represented as mean ± SD. **(B)** Representative maximum intensity projections of myosin II heavy chain (left) and fibronectin (middle) on mLCNs after 5 days of differentiation. Scale bars = 200 µm. Corresponding frequency distribution plot (right) of myosin and fibronectin alignment. Data are represented as mean ± SD. **(C, D)** Frequency distribution plots of myosin and **(C)** Laminin, or **(D)** Collagen IV alignment after 5 days of differentiation. Data are represented as mean ± SD.

Contractile cells are known to locally align their ECM, which in turn supports greater forces and amplifying anisotropic tension through the ECM that can align cells over large distances.[40,41] In one report on these reciprocal cell-ECM dynamics, increasing fibroblast contractility exceeded the tensile strength of the nascent ECM, driving the eventual formation of contractile instabilities.[42] Myotubes are known to become highly contractile, with a nascent myotube doublet exerting more than twice the summed force of two unfused myoblasts.[43] Towards the end of our live-cell imaging experiments, we observed similar instabilities as highly contractile myotubes overpowered the ECM and created holes in the culture **(Figure 2E; Video S2)**. These holes developed along the axis of the cellular flows, with a slight CW offset to the nematic director in concordance with the offset myotube alignment in **Figure S3**. While the ECM is known to be viscoelastic on short time scales, it also exhibits plasticity when forces are applied over longer time scales.[44] Therefore, we hypothesized that increasingly contractile myotubes exert coordinated, persistent forces on the ECM causing plastic deformation and reorienting/displacing myotubes attached to and enmeshed within it.

Accordingly, we tested whether we could prevent global myotube-ECM alignment on mLCN-1.0 by disrupting the positive feedback loop between increasingly contractile myotubes and the viscoplastic ECM. We treated C2C12s on mLCNs with the non-muscle myosin II inhibitor blebbistatin daily for three days, beginning at confluence (t_a_ = 0 hr) and imaging every 3 hours **(Video S3)**. Shortly after the addition of 10 µM blebbistatin, C2C12 order decreased from 0.45 to 0.3, with a corresponding decrease in ξ_nn_ from 600 µm to 450 µm **(Figure 4A)**. Blebbistatin had a delayed effect on ξ_vv,_ which continued to increase in tandem with the control samples until t_a_ = 42 hr when it diverged and returned to the baseline velocity correlation length of ∼240 µm. We attribute this delayed effect to the moderate ECM alignment found at confluence, which may provide mechanical memory for cellular migration.[45] Meanwhile, C2C12 order, ξ_nn_, and ξ_vv_ increased as expected for samples not treated with blebbistatin **(Figure 4A)**. While the characteristic increase in ξ_vv_ with decreasing speed over time was again observed on uninhibited samples, the opposite trend was seen with blebbistatin treatment **(Figure 4B)**.

**Figure 4.**
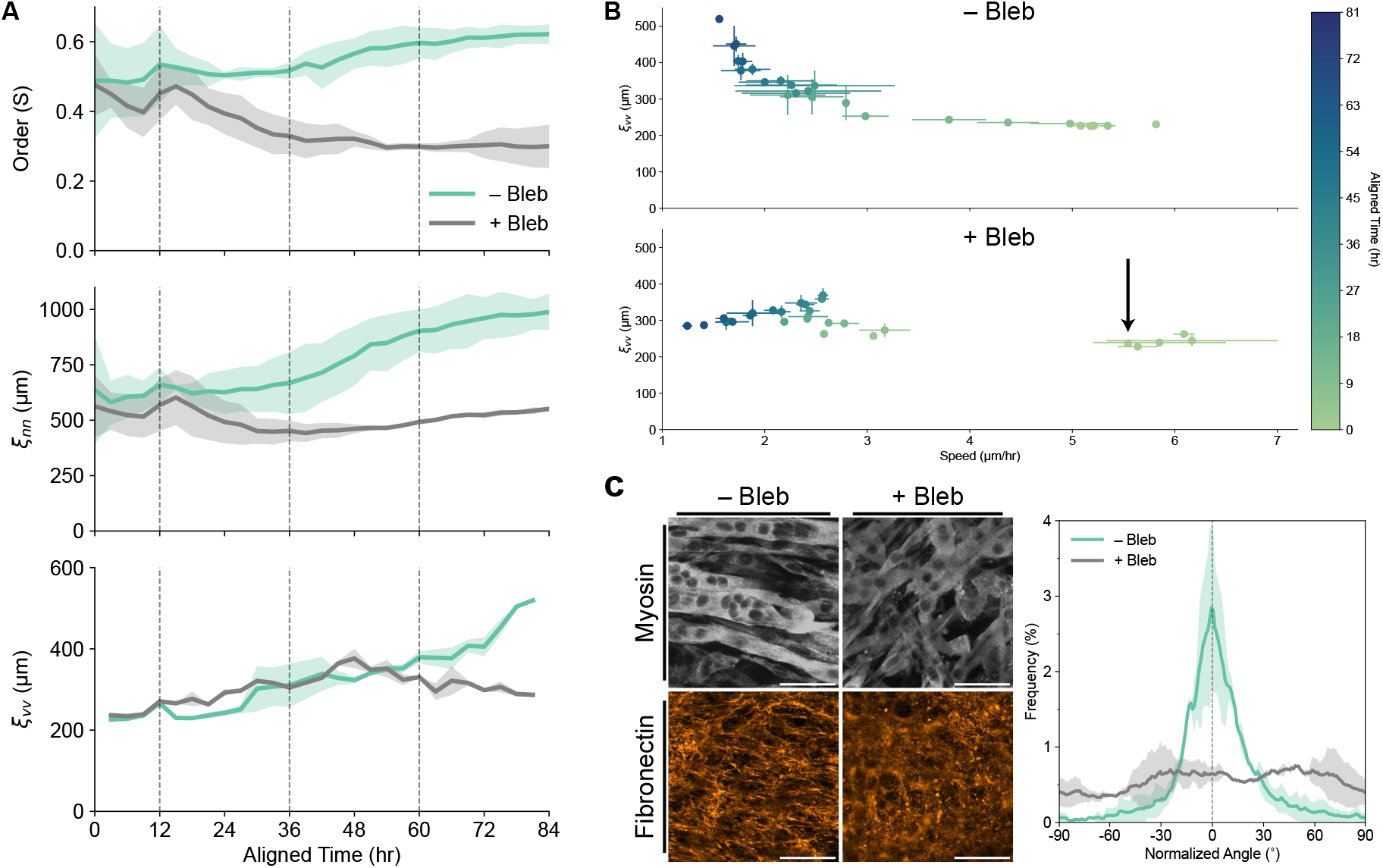
Reciprocal cell-ECM dynamics mediate collective cellular flows on mLCNs. (A) Top to bottom: Temporal evolution of orientation-order parameter (S), nematic correlation length (µm), and velocity correlation length (µm) on mLCNs ± blebbistatin treatment. Dashed lines indicate timing of daily media changes, with the first addition of blebbistatin occurring at t_a_ = 12 hr. Data are represented as mean ± SEM. **(B)** Velocity correlation length (µm) as a function of cellular speed (µm/hr) on mLCNs ± blebbistatin treatment. Data are represented as mean ± SEM. Arrow indicates first addition of blebbistatin. **(C)** Representative maximum intensity projections (left) of myosin II heavy chain (top) and fibronectin (bottom) after 3.5 days of differentiation on mLCNs ± blebbistatin treatment. Scale bars = 50 µm. Corresponding frequency distribution plot (right) of fibronectin alignment after 3.5 days of differentiation on mLCNs ± blebbistatin treatment. Data are represented as mean ± SD. See also **Video S3**.

Finally, we immunostained these samples for myosin and fibronectin to assess the effect of disrupting cellular contractility on myotube fusion and ECM organization. Although myoblasts were still capable of differentiation and fusion in the presence of blebbistatin (confirmed by positive myosin II heavy chain staining), the resulting myotubes were shorter and disorganized **(Figure 4C)**. We again found dense, aligned fibronectin on untreated samples, whereas on blebbistatin-treated samples, fibronectin staining was sparse and completely isotropic **(Figure 4C)**. These results demonstrate that coordinated supracellular contractility, which was disrupted with blebbistatin, is necessary for the emergence of contractile cell-ECM flows that drive global myotube-ECM alignment with the nematic director of mLCNs.

## Discussion

### Stiffness anisotropy is a mechanobiological cue for collective cell-ECM alignment

Engineered biomaterials are increasingly used to study and direct biophysical cellular processes that give rise to the organized multicellular morphologies found in tissues and organs.[46] This work provides an integrated analysis of how anisotropy can be transmitted and amplified across length scales from the liquid crystalline mesogen (∼10 Å) of mLCNs to skeletal muscle myotube cultures spanning several millimeters. Molecular anisotropy of LC mesogens is translated to mechanical anisotropy in the resulting polymer network, which serves as a mechanobiological cue to C2C12s enabling them to coordinate contractility in the stiffest direction (i.e., with the nematic director) resulting in global myotube-ECM alignment on the mLCN substrate. In the absence of anisotropy (i.e., when LCNs are polymerized in the isotropic state), myotubes do not develop long-range order, with 2D order and ξ_nn_ similar to that of myotubes grown on glass coverslips **(Figures 1 and S3)**. Since the mLCNs had minimal topographic features, well below the requirement for contact guidance, we conclude that C2C12s are able to collectively sense the stiffness anisotropy of mLCNs.

Our use of an aza-Michael addition chemistry enabled us to systematically investigate how the degree of stiffness anisotropy in three different mLCNs impacts myotube alignment without any confounding effects from topographic contact guidance. Although mLCN-0.5 displayed the greatest stiffness anisotropy ratio (E_||_ / E_1−_), myotube order and ξ_nn_ was higher on mLCN-0.75 and mLCN-1.0 **(Figures 1 and S3)**, suggesting that myoblasts are sensing the orthogonal difference in elastic modulus (E_||_ – E_1−_). Thrivikraman *et al.* also found that the degree of fibroblast alignment on anisotropic fibrin gels scaled with the orthogonal difference in elastic modulus.[47] However, the difference in overall stiffness between mLCN-0.5 and the two stiffer mLCNs cannot be overlooked and may contribute to the observed differences. We also found that myotube order and ξ_nn_ was significantly higher on mLCN-1.0 compared to mLCN-0.75. As the only difference between these formulations is a greater degree of stiffness anisotropy for mLCN-1.0, this comparison demonstrates that collective cellular alignment scales with the degree of stiffness anisotropy.

Through live-cell imaging and advanced image analysis, we identified three distinct temporal phases through which C2C12s interact with the mLCN substrate, neighboring cells, and nascently deposited ECM over increasing length scales to generate long-range order on mLCNs: (i) cellular crowding, (ii) collective myoblast polarization, and (iii) contractile cell-ECM flows mediated by reciprocal cell-ECM dynamics **(Figure 5)**. Across all three mLCN formulations, stiffness anisotropy had no effect on myoblast polarization or migration before confluence **(Figure S2)**. This result was not unexpected, given multiple reports of single cells being unresponsive to stiffness anisotropy on substrates with high baseline stiffness.[6,7,48] However, myoblasts exhibited enhanced sensitivity to stiffness anisotropy with increasing cell density, similar to the collective durotaxis of cell monolayers cultured on hydrogels with linear stiffness gradients that were undetectable by single cells.[4] While epithelial monolayers undergo jamming transitions upon reaching confluence,[36,49] we found that C2C12 myoblasts continue proliferating after reaching confluence and form a distinct multilayered culture **(Figures 2A–C)**.[37] During the proliferative phase, myoblast migration speed decreased and order rose as cell crowding induced nematic packing,[50] leading to local domains of alignment highlighted by the presence of topological defects.[51]

**Figure 5.**
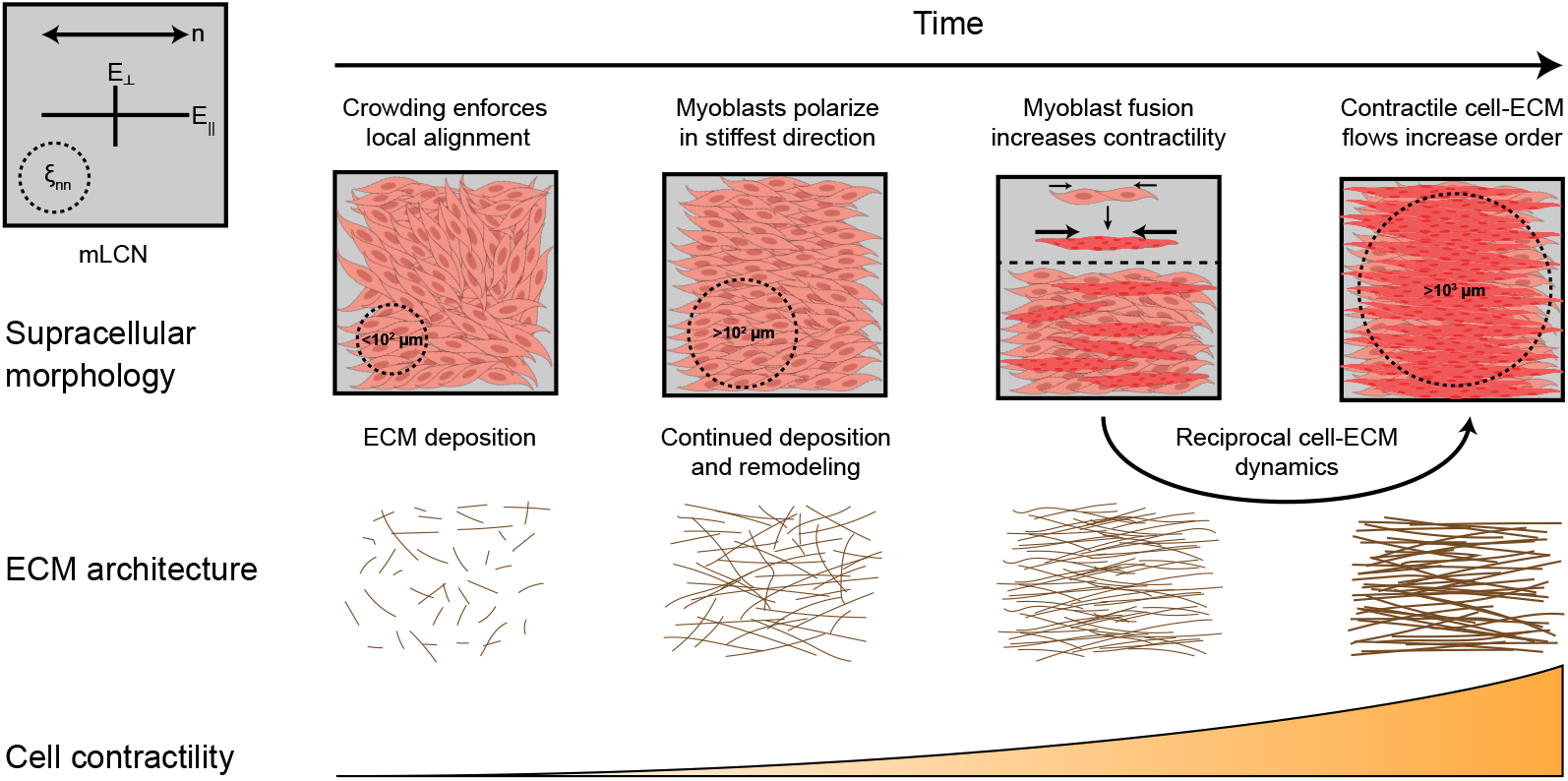
Spatiotemporal evolution of collective cellular dynamics that drive C2C12 myoblast and myotube alignment with the nematic director of mLCNs. Created with BioRender.com.

Myoblast order on mLCNs was higher at confluence compared to iLCNs **(Figure 2C)**, suggesting that myoblasts on mLCNs begin polarizing in the stiffer direction once a critical cell density threshold is reached. Collective polarization in migrating cellular monolayers has been observed previously, whereby cells reorient in the direction of greatest normal stress to minimize intercellular shear stress in a process termed plithotaxis.[50] However, we observed myoblast reorientation before the emergence of collective cellular flows on mLCNs. Still, these processes may be related. As the network of cell-cell and cell-substrate contacts mature, myoblasts coordinate actomyosin contractility in the direction that they can generate the greatest stress (i.e., the nematic director of the mLCN) and polarize accordingly.

ECM deposition and remodeling before and during collective myoblast polarization resulted in a moderately-aligned ECM architecture that both supports increased contractile forces and propagates these forces across large distances through reciprocal cell-ECM dynamics.[41,42] Shortly after proliferation was arrested, myoblasts began fusing into nascent myotubes capable of generating exponentially greater contractile forces. These forces resulted in plastic deformation of the cell-secreted ECM and highly correlated collective cellular flows, as any myotubes embedded within and adhered to the matrix flowed with it. PIV analysis revealed increasing correlation length of these flows only on mLCNs **(Figure 2C)**, likely due to a combination of preferential anisotropic fusion along the axis of collective myoblast polarization, moderate ECM alignment before fusion, and the continued influence of substrate stiffness anisotropy on collective dynamics. These long-range contractile cell-ECM flows on mLCNs resulted in annihilation of most topological defects in the myotube cultures, reorienting nearly all myotubes in the direction of greatest stiffness **(Figure 2E; Video S2)**. This positive feedback loop between increasing cellular contractility and ECM alignment ultimately results in remarkable global ordering of C2C12 myotubes on mLCNs across several millimeters. As further evidence of these processes, inhibiting C2C12 contractility with blebbistatin disrupted the coordination of actomyosin contractility on mLCNs. Blebbistatin impacted both collective myoblast polarization and contractile cell-ECM flows, which ultimately abrogated long-range myotube and ECM alignment **(Figure 4)**.

### Stiffness anisotropy organizes active stresses to guide tissue morphogenesis

These findings offer a unique comparison to the biophysical evolution of other collective cellular dynamics such as the aforementioned jamming transitions in epithelial monolayers, which have been shown to be highly sensitive to the relative strength of cell-cell and cell-matrix contacts.[36] For example, where we found that C2C12s with weak cell-cell adhesions exhibit increasing velocity correlation length as cell density increases and velocity correspondingly decreases **(Figure 2D)**, the exact opposite was observed in epithelial cells with strong cell-cell adhesions.[36] Transient blebbistatin treatment in otherwise glassy epithelial monolayers increased long-range velocity correlation.[52] Meanwhile, when C2C12s were treated with blebbistatin, we observed decreasing velocity correlation length with decreasing speed over time **(Figure 4B)**.

In a more direct comparison, fibroblast alignment with the nematic director of mLCNs was found to be driven primarily by directed cell division along the substrate’s nematic axis and associated extensile stresses that caused annihilation of topological defects in the monolayer.[15] An earlier study also found that epithelial cells exhibited collective extensile behavior, while mesenchymal cells collectively behaved as a contractile system.[53] When cell-cell adhesions were weakened by knocking out E-cadherin in epithelial cells, their collective behavior switched from extensile to contractile with a concomitant increase in cell-substrate traction forces. Conversely, when the ability of E-cadherin knock-out cells to generate substantial traction forces was reduced, either by culture on a soft substrate or when contractility was inhibited with blebbistatin, the cellular collectives switched back to extensile behavior.[53] Thus, the interplay between cell-cell and cell-substrate (or cell-matrix) interactions dictates the nature of active stresses that emerge once cells become confluent and behave as a collective. In our system, C2C12 collective dynamics are dominated by contractile mechanisms associated with cell-substrate (e.g., collective polarization in response to stiffness anisotropy) and cell-matrix (e.g., reciprocal cell-ECM dynamics) interactions.

These comparisons highlight the influences of cell identity on active nematic behaviors that guide tissue morphogenesis. Crucially, it appears that stiffness anisotropy of mLCNs can establish anisotropy of both contractile and extensile stresses to drive collective cellular alignment independent of cell type. The development of nematic-like alignment in cellular collectives may be necessary for the emergence of higher-level nematic cellular interactions, such as the cholesteric-like chiral layering of aligned C2C12 myotubes observed on both mLCNs and NanoSurface substrates **(Figure S4)**. Taken together, our multi-scale analysis and other studies of collective cellular dynamics reveal the mechanisms by which cell populations are controlled by biophysical interactions to orchestrate distinct coordinated behaviors in development and regeneration.

### Skeletal muscle alignment by emergent collective cellular dynamics

In the context of skeletal muscle tissue, our study identifies fundamental collective cellular mechanisms for myotube alignment that may be relevant to skeletal muscle development and regeneration *in vivo*. Our results suggest that the mechanical anisotropy (e.g., of nascent ECM during development or basal lamina remnants following injury) is an effective cue for collective myoblast polarization prior to fusion. Myoblasts must also secrete a provisional ECM to support (re)generation of fully functional tissue.[19,39,54] As myoblasts fuse and develop contractile stresses in this developing matrix, the emergence of reciprocal cell-ECM dynamics would further align myotubes and their ECM to support the large forces associated with muscle contraction and promote maturation of fully functional myofibers. In support of this, gradients of anisotropic stresses that are contractile anteriorly and extensile posteriorly were shown to be responsible for shaping the tissue-level chevron patterns characteristic of zebrafish myotomes.[55] As interest continues to grow in developmental biophysics, dynamic interactions between populations of cells and their underlying ECM mechanics might be subject to future study.

While molecular and genetic events inside a single cell are required to initiate muscle development and regeneration, the emergence of collective cellular behaviors are equally important. Investigations of these collective cellular dynamics *in vitro* necessitate the continued development and use of engineered biomaterials with tunable mechanical properties, highlighted by our use of LCNs in the present study. LCNs are uniquely amenable to spatial patterning of nematic alignment and thus mechanical anisotropy, opening new doors to investigate the interplay between substrate mechanics and the active stresses that arise in cellular collectives and guide tissue shape and organization.

## Limitations of the study

It is important to highlight that the overall moduli (MPa–GPa) of the materials studied here, as well as their large (MPa) orthogonal differences in moduli, are much greater than that of most soft tissues, including muscle.[56] Furthermore, these surface-aligned mLCNs restrict studies of collective cellular dynamics to the 2D cell-material interface. Future studies will focus on the development of innovative liquid crystalline polymer networks capable of 3D cell encapsulation with physiologically-relevant mechanical properties to more accurately understand how myoblasts collectively respond to stiffness anisotropy *in vivo*. Oriented, tunable biomaterials that more faithfully recapitulate the anisotropic mechanical properties of tissues and their ECM will enable the field to probe intrinsic biophysical processes that are essential to developmental and regenerative processes. Further, the ability to understand and harness these dynamics to program multicellular morphology and function holds great promise for the regenerative biology, medicine, and engineering communities.

## Acknowledgements

The authors gratefully acknowledge funding from NIH F30HL164047 (N.P.S), DARPA W911NF- 19-2-0024 (B.E.K., B.R.N), CU Boulder Biological Sciences Initiative (G.K.H.), Army Research Office W911Nf-19-1-0349 (K.M.H., T.J.W.) and NIH R01 DE016523 (K.A.G., K.S.A). F.W.D and R.M.K. were supported by the Center for Integrated Nanotechnologies, a Department of Energy office of Basic Energy Sciences user facility. Sandia National Laboratories is a multimission laboratory managed and operated by National Technology & Engineering Solutions of Sandia, LLC, a wholly owned subsidiary of Honeywell International Inc., for the U.S. Department of Energy’s National Nuclear Security Administration under contract DE- NA0003525. This paper describes objective technical results and analysis. Any subjective views or opinions that might be expressed in the paper do not necessarily represent the views of the U.S. Department of Energy or the United States Government. We thank Dr. Margaret Prendergast for sharing and assisting in adapting her circular statistics R script. We thank Drs. Qiyan Mao and Frank Schnorrer for sharing and assisting in adapting their R scripts to calculate 2D order and correlation lengths. We also thank Dr. Joselle McCracken, Connor Miksch, Dr. Matthew Davidson, Dr. Jason Silver, and Dr. Joseph Dragavon for experimental assistance and/or helpful discussions. Some imaging work was performed at the BioFrontiers Institute Advanced Light Microscopy Core (RRID: SCR_018302) on a Nikon A1R confocal laser scanning microscope supported by NIST-CU Cooperative Agreement award number 70NANB15H226.

## Author Contributions

Conceptualization, N.P.S, B.E.K., K.M.H., T.J.W., and K.S.A.; Methodology, N.P.S., B.E.K., K.M.H., K.A.G., R.M.K., and F.W.D.; Formal Analysis, N.P.S., B.E.K., K.M.H., B.R.N., G.K.H., R.M.K., and F.W.D.; Investigation, N.P.S., B.E.K., K.M.H., K.A.G., R.M.K., and F.W.D.; Writing – Original Draft, N.P.S. and B.E.K.; Writing – Review & Editing, N.P.S., B.E.K., K.M.H., B.R.N., G.K.H., K.A.G., R.M.K., F.W.D., T.J.W., and K.S.A.; Visualization, N.P.S. and B.E.K.; Resources, F.W.D., T.J.W. and K.S.A.; Supervision, T.J.W. and K.S.A.; Funding Acquisition, T.J.W. and K.S.A.

## Methods

### RESOURCE AVAILABILITY

#### Lead Contact

Further information and requests for resources and reagents should be directed to and will be fulfilled by the lead contact, Kristi S. Anseth.

#### Materials Availability

This study did not generate new unique reagents or materials.

#### Data and Code Availability

All data reported in this paper will be shared by the lead contact upon request.

All original code will be deposited in a publicly available repository before the time of publication. Any additional information required to reanalyze the data reported in this paper is available from the lead contact upon request.

### EXPERIMENTAL MODEL AND STUDY PARTICIPANT DETAILS

#### Cell Lines

C2C12 murine myoblasts (CRL-1772, ATCC) were passaged and cultured on TCPS (Grenier) for use up to passage 15. The sex of this cell line is unknown. C2C12s were cultured at 37 °C and 5% CO_2_ in sterile-filtered growth media, which consisted of High Glucose Dulbecco’s modified Eagle’s Medium (DMEM, Gibco) supplemented with 20% FBS (Gibco), 1x GlutaMAX (Gibco), 1 mM sodium pyruvate (Gibco), 50 U/mL penicillin (Gibco), 50 µg/mL streptomycin (Gibco), and 1 µg/mL amphotericin B (Gibco).

### METHOD DETAILS

#### LCN Synthesis

Liquid crystalline monomer 1,4-Bis-[4-(6-acryloyloxyhexyloxy)benzoyloxy]-2-methylbenzene (C6M, Wilshire Technologies), hexylamine (Sigma-Aldrich), thermal inhibitor 2,6-Di-*tert*-butyl- 4-methylphenol (BHT, Sigma-Aldrich), and photoinitiator Irgacure-651 (I-651, Sigma-Aldrich) were used. C6M, 2 wt% I-651, and 1 wt% BHT were added to a glass vial, melted at 100 °C and vortexed to ensure homogeneity. Hexylamine was added at 85 °C to achieve the final acrylate molar fraction of 0.5 and 0.75 in LCN-0.5 and LCN-0.75, respectively. The mixture was again vortexed and drawn by capillary action into 50 µm thick alignment cells at 85 °C. The alignment cells were fabricated using Elvamide-coated glass slides that were rubbed with velvet and adhered in anti-parallel arrangement with a mixture of optical adhesive and 50 µm spacers. The filled alignment cells were held for 18 hours at 75 °C to facilitate aza-Michael addition within the nematic phase. Oligomers were subsequently photopolymerized within the alignment cell at 75 °C with 15 minutes of UV (365 nm) light at 10 mW/cm^2^. In the case of LCN-1.0, monomers were melted, mixed, and drawn into cells as described above, but proceeded directly to photopolymerization as no hexylamine was added. The cells were soaked in water and carefully opened to remove the polymerized films. Isotropic LCNs were synthesized in the same manner but polymerized in the isotropic state at 135 °C in Elvamide-coated glass cells that were not rubbed.

#### Polarized Optical Microscopy

Molecular alignment of mLCNs, or lack thereof for iLCNs, was confirmed by the presence or absence of birefringence using a Nikon Eclipse Ci-Pol with a 5x objective. LCNs were observed at 0° and 45° rotation between crossed polarizers.

#### Tensile Testing

mLCNs were cut into strips parallel and perpendicular to the nematic director. Tensile tests were performed using an RSA-G2 DMA (TA Instruments) at a constant rate of 5% strain/min until failure. The elastic modulus values reported in **Figure 1B** are calculated from the slope of the initial linear stress/strain regime (See also **Figure S1E**). n = 4 (mLCN-0.5_||_), 2 (mLCN-0.5_1−_), 6 (mLCN-0.75_||_), 3 (mLCN-0.75_1−_), 2 (mLCN-1.0_||_), 3 (mLCN-1.0_1−_).

#### Contact Angle Measurements

Contact angle experiments were performed using a custom contact angle setup that included a syringe pump (Aitoserlea) and high-speed camera (Plugable). While imaging continuously with the camera, a small (∼4 µL) droplet of deionized water was extruded through a 32 gauge needle (Nordson) driven by the syringe pump until the surface tension released the droplet onto the LCN. The contact angle was determined using the contact angle plugin for ImageJ assuming a spherical droplet. n = 3 independent mLCNs for each formulation.

#### Atomic Force Microscopy

Surface roughness measurements were conducted on an Asylum MFP-3D atomic force microscope (AFM). LCNs were mounted on stainless-steel AFM pucks using double-sided tape. Rectangular Si cantilevers with reflective Al coatings on the detector side were utilized to enhance spatial resolution and laser reflectivity (Nanosensors PPP-NCHR cantilevers with a nominal spring constant *k*_c_ = 42 N/m and tip radius *R* = 7 nm). Intermittent-contact mode images of surface heights *z* were collected at a scan rate of 1 Hz. The images were taken with a 5 μm × 5 μm scan area and 512 scan points and lines, which translated to a ≈ 10 nm pixel size. Subsequently, the images were processed with first-order plane fit and flattening routines to account for sample tilt. Cross-sectional profiles of the *z* maps were amassed to enable quantitative comparisons between the various films; the profiles presented in **Figure S1G** were typical of data from other regions in each image. Similarly, the root mean square (rms) roughness values were reported for each image to evaluate any relationship between average topography and surface response. n = 1 mLCN for each formulation.

#### Substrate Preparation for Cell Culture

LCNs were cut into nearly-square rectangles (∼5x4 mm) with either the long or short axis corresponding to the nematic director, depending on the experiment. LCNs were sterilized with 70% Ethanol/H_2_O, placed into a 24-well plate, and washed with sterile PBS. 12 mm glass coverslips were sterilized with 70% Ethanol/H_2_O, placed into a 24-well plate, and washed with sterile PBS. 35 mm NanoSurface dishes (Curi Bio) were sterilized with 70% Ethanol/H_2_O and washed with sterile PBS. 300 µL of 0.1% gelatin in water (StemCell Technologies) was added to the coverslips and NanoSurface dishes, which were incubated for >30 minutes at 37 °C to allow the gelatin to cure. Excess gelatin solution was removed from coverslips and NanoSurface dishes before cell seeding.

#### C2C12 Seeding, Growth, and Differentiation

C2C12 myoblasts were trypsinized from TCPS and resuspended in fresh growth media. 1 mL of growth media containing 5,000 cells/cm^2^ were added to the wells with LCNs, gelatin-coated coverslips, or NanoSurface dishes. Cells were allowed to adhere to LCNs for 2 hours, at which time they were transferred to new wells with 1 mL growth media to ensure cells were only present on the LCNs. Cells were cultured in growth media for 72 hours until ∼70–80% confluent. At this point, growth media was replaced with sterile-filtered differentiation media to induce differentiation via serum-starvation. Differentiation media consisted of High Glucose Dulbecco’s modified Eagle’s Medium (DMEM, Gibco) supplemented with 5% horse serum (Gibco), 50 U/mL penicillin (Gibco), 1x GlutaMAX (Gibco), 1 mM sodium pyruvate (Gibco), 50 µg/mL streptomycin (Gibco), and 1x ITS Supplement (R&D Systems). Differentiation media was replaced daily for up to 5 days, at which point cells were fixed.

#### Immunofluorescence Staining

All steps were carried out at room temperature. Samples were washed once with PBS and fixed with 4% paraformaldehyde (Electron Microscopy Science) diluted in PBS for 10 minutes. Samples were washed with PBS three times for 5 minutes each. Samples then were permeabilized and blocked with 0.1% Triton-X in PBS containing 3% BSA (Sigma-Aldrich) for 30 minutes. Samples were incubated with primary antibodies diluted in the same buffer for 1 hour. The following primary antibodies were used: MF-20 (1:250, eBioscience, 14-6503-83), Fibronectin (1:500, Abcam, ab2413), Laminin (1:500, Abcam, ab11575), and Collagen IV (1:500, Abcam, ab19808). After washing with PBS three times for 5 minutes each, samples were incubated with secondary antibodies in the same buffer as primary antibodies for 1 hour in the dark. The following secondary antibodies were used: Alexa Fluor 488 goat anti-mouse (1:250, Invitrogen, A-11001), Alexa-Fluor 647 goat anti-rabbit (1:500, Invitrogen, A-21245), Rhodamine Phalloidin (1:500, Cytoskeleton, PHDR1), Alexa Fluor 488 Phalloidin (1:500, Invitrogen, A12379), and DAPI (1 µg/mL, Sigma-Aldrich, 10236276001). Samples were washed with PBS three times for 5 minutes each before imaging.

#### Imaging

Fixed samples were imaged on multiple microscopes including a Zeiss LSM710 laser scanning confocal equipped with 20x/1.0 NA water objective, a Nikon A1R laser scanning confocal equipped with 10x/0.45 NA Air or 20x/0.95 NA LWD water objective, or a Nikon Ti-E microscope equipped with 10x/0.45 NA and 20x/0.75 NA air objective, an Okolab environmental chamber and a CREST X-Light V2 spinning disk system.

#### Live-Cell Imaging

C2C12 myoblasts were seeded on LCNs as described above. Once cells were adhered, SiR-Actin (1:1000, Cytoskeleton, CY-SC001) and/or SPY-555-DNA (1:2000, Cytoskeleton, CY-SC201) was added to each well. Samples were imaged every 1/3, 1, 3, or 6 hours using a Nikon Ti-E microscope equipped with 10x/0.45 NA or 20x/0.75 NA air objective, an Okolab environmental chamber (37 °C, 5% CO_2_, 95% relative humidity) and a CREST X-Light V2 spinning disk system.

#### Image Analysis

All image analysis was conducted in ImageJ except for PIV analysis conducted in MATLAB. Maximum intensity projections were used for most analysis. For myotubes cultured on mLCNs, the images were rotated such that the nematic director of the mLCN was parallel to the horizontal (0°) axis. For myotubes cultured on coverslips or iLCNs, the images were not rotated. For myotubes cultured on NanoSurface dishes, images were rotated such that the nanopatterned grooves were parallel to the horizontal (0°) axis. All images were cropped to exclude cells within 500 µm of the edges of the LCN, coverslips, and NanoSurface dishes due to contact-guidance effects of free edges. For the single cell analysis in **Figure S2**, images were thresholded, segmented, and ellipses were fit to individual cells. The number of cells, area, aspect ratio, and orientation was quantified with the Analyze Particles tool in ImageJ. Orientation counts were binned in 20° increments, converted to frequencies, and averaged across replicates. Cell density, area, and aspect ratio parameters were averaged for all cells in a single sample and timepoint and then averaged across replicates. n = 3 independent samples for each substrate at each timepoint. For the intensity frequency plots in **Figure S4**, composite images were resliced to XZ projections without interpolation and summed across all Y. Then, average intensities of each z-slice were taken across all X. Intensity values were converted to frequencies, averaged across independent samples, and displayed as mean ± SD. For the axis of myoblast division in **Figure S5**, individual frames of SPY-555-DNA-stained nuclei were examined and a line was manually drawn between the center of two nuclei of dividing cells. The orientation of these lines were taken with respect to the horizontal axis (i.e., nematic director), counts were binned in 20° increments, converted to frequencies, and averaged across replicates.

#### OrientationJ Analysis

OrientationJ analysis[33] was performed on each image with σ = 10 or 20 pixels (for 10x and 20x objective, respectively) for myotube analysis and σ = 1 or 2 pixels (for 10x and 20x objective, respectively) for ECM analysis, with minimum coherency set to 25%. When the confocal microscope was used to acquire images with greater resolution **(Figures 4C and S4)**, σ was set to the approximate width of myotubes or ECM fibrils in pixel units. Counts from the OrientationJ results were converted to frequencies. For ECM alignment analysis in **Figures 3B–D and S4**, the orientation frequency distribution was shifted such that the angle of maximum myotube alignment for each sample was 0°. This was done due to the variation in the degree of myotube offset to the nematic director (**See Supplementary Note 1, Figure S6**). The frequency distribution of ECM orientation was then shifted by the same degree. When this strategy was employed, the x-axis reads ‘Normalized Angle (°)’. Frequencies were then averaged and displayed as mean ± SD. For the chiral layer analysis in **Figure S4**, maximum intensity projections of each layer were used and OrientationJ analysis was performed on each image as described above.

#### Cell Density and Timeseries Alignment

Timeseries images of SPY-555-DNA-stained nuclei were imported into ImageJ and nuclei were detected using the StarDist plugin (model: versatile (fluorescent nuclei), probability threshold = 0.65, nmsThreshold = 0.55).[57,58] The spots were exported to the ROI manager and the number of nuclei was determined by running the Analyze Particles tool. Cell density was calculated by dividing the number of cells at each timepoint by the area in mm^2^. Due to the dependence of cellular alignment on cell density and small experimental variation in initial cell seeding density, the start time of every timeseries was adjusted such that the maximum cellular density was achieved at t_a_ ∼ 36 hr, which corresponded to cells achieving confluence at t_a_ ∼ 0 hr. All other analyses (order, spatial disorder, ξ_vv_, speed) reported in **Figures 2C and 2D** were adjusted to the start times determined by this method. This resulted in some timepoints having just one value for cell density, order, spatial disorder, *ξ_vv_*, and speed and thus no standard error is reported for those points. Live-cell imaging for **Figures 2E and 2F** ended after 5 days of differentiation and thus the final timepoint was set as t_a_ = 120 hr. Live-cell imaging for **Figure 4** started once cells reached confluence and thus at t_a_ = 0 hr.

#### Migration Analysis

Timeseries images of SPY-555-DNA-stained nuclei were locally enhanced with the contrast-limited adaptive histogram equalization (CLAHE) tool (blocksize: 100; histogram bins: 256; maximum slope: 3) and imported into the ImageJ plugin TrackMate where nuclei were detected using the StarDist tracker.[57,58] Nuclei with diameter larger than 30 µm or smaller than 6 µm were excluded. Tracks were created using the Simple LAP Tracker with default settings. Spots and Tracks were exported to a MotilityLab spreadsheet and imported into MATLAB where a custom script calculated mean speed and angle of migration. The normalized cellular migration within ± 10° of the nematic director was calculated by first obtaining the angle of migration of each cell at each timepoint, then calculating the summed frequency of cells falling into 2 bins (-10° – 0°, 0° – 10°) and dividing by the frequency for isotropic migration (2 bins out of 18 total = 11.111%). Mean migration speed and normalized migration within ± 10° of the nematic director as a function of cell density was obtained by averaging linear interpolations of individual migration analyses with increments of 10 cells/mm^2^. n = 12 non-overlapping fields of view across 3 independent mLCNs.

#### PIV Analysis

PIV analysis was conducted using the PIVlab version 2.61 software plugin for MATLAB. For all PIV analysis, image pre-processing parameters included CLAHE: 64 pixels and auto contrast stretch. PIV settings for **Figures 2C and 2D** were: Pass 1: 300 pixel interrogation area / 150 pixel step size, Pass 2: 150 pixel interrogation area / 75 pixel step size. PIV settings for **Figure 2F** were: Pass 1: 600 pixel interrogation area / 300 pixel step size, Pass 2: 300 pixel interrogation area / 150 pixel step size, Pass 3: 150 pixel interrogation area / 75 pixel step size. PIV settings for **Figures 4A and 4B** were: Pass 1: 1000 pixel interrogation area / 500 pixel step size, Pass 2: 500 pixel interrogation area / 250 pixel step size, Pass 3: 250 pixel interrogation area / 125 pixel step size. Vectors were smoothed at the lowest setting and aberrant vectors at the frame boundaries were manually rejected to remove artifacts. Velocity magnitudes were averaged across each frame and then across independent samples and converted to µm/hr. After media changes, the field of view tended to shift slightly resulting in artificially high autocorrelation at certain timepoints. While others have subtracted the mean velocity at each timepoint to eliminate this artifact, this is only applicable for isotropic systems.[36] In our system the flows are polarized and thus we chose to remove those artifactual datapoints from our analysis altogether. These frames are dimmed in **Videos S1–S3**, which display autoscaled vectors overlaid on a heatmap of velocity magnitude ranging from 0 – 5 µm/hr **(Video S1)**, 0 – 8 µm/hr **(Video S2)**, and 0 – 7 µm/hr **(Video S3)**.

#### Line Integral Convolution

OrientationJ VectorField was run on images of 2500x2500 pixels with σ = 10 pixels and grid size = 5 pixels. Resulting .csv files were imported into python where the ‘lic’ package (version 0.4.5) was used to create the images in **Video S2** with the ‘length’ variable set to 20.

### QUANTIFICATION AND STATISTICAL ANALYSIS

#### 2D Orientation-Order Parameter (S)

Images were cropped into squares with pixel size a multiple of 100 (e.g., 1300 x 1300 pixels) and OrientationJ VectorField was run with σ = 10 or 20 pixels and grid size = 100 or 50 pixels (for 10x and 20x objective, respectively). 2D orientation-order parameter and nematic correlation length were calculated using custom R scripts adapted from Mao et al.[28] Order was calculated using the orientation of each vector with the following equation[59] where θ represents the difference between the angle of an individual vector and the mean of all angles in the image and the brackets < > denote an average over grid points.

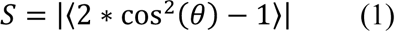

This equation provides the 2D orientation-order parameter in a range from 0 to 1, where 0 is completely isotropic and 1 is perfectly order. For global order calculations, S was calculated over all angles in the image. For spatial disorder and local order calculations, S was calculated for 4x4 grids of vectors at each center position. Spatial disorder was calculated as the percentage of grids with an orientation-order parameter below a threshold of 0.5. For the grayscale order heatmap in **Video S2**, local orientation-order parameter (0–1) at each position was scaled to 8-bit grayscale values (0–255), imported into ImageJ as a text image and saved as .tiff file.

#### Nematic and Velocity Correlation Length

Nematic correlation length (in pixel units) was taken as the X-intercept of the least-squares regression of the first 10 points in the spatial autocorrelation function of the vector field output from OrientationJ:

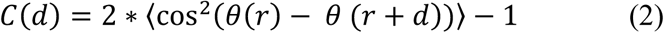

Where θ(r) is the local orientation angle of the vector field at grid position r, d is the separation vector to a second grid position r+d at distance |d|=d, and brackets < > denote an average over all pairs of grid points with the same distance d. Nematic correlation length was converted to µm by multiplying it with the pixel size (µm/pixel) of each image. For velocity correlation length calculations, vector angles from PIVlab were passed to a custom MATLAB script to format the data for import into R. The same script and equation (2) used to calculate the nematic correlation length was used to calculate the velocity correlation length, again taking the X-intercept of the least-squares regression of the first 10 points in the spatial autocorrelation function and multiplying it with the pixel size (µm/pixel) of each image to obtain the velocity correlation length in µm.

#### Statistical Analysis

Data are shown as the mean ± standard deviation or standard error, unless otherwise indicated in the figure legend. All n values refer to biologically independent samples and significance was claimed at \**p*<0.05, \*\**p*<0.01, \*\*\**p*<0.001, \*\*\*\**p*<0.0001. Sample size for statistical comparisons was n ≥ 3 unless otherwise indicated. Circular statistics were performed using a custom script with the package “circular” in R adapted from Davidson et al.[60] The Watson-Wheeler test for homogeneity was used to compare myotube alignment on monodomain and isotropic LCNs of the same formulation and timepoint. GraphPad Prism 9 was used for the remainder of statistical analysis. Statistical comparisons between two experimental groups were assessed via one-tailed or two-tailed unpaired Student’s *t*-test with Welch’s correction. One-tailed *t*-test was performed to compare orientation-order parameter and nematic correlation length of myotubes cultured on monodomain vs. isotropic LCNs as the hypothesis was that cells would align better on monodomain LCNs. Two-tailed *t*-tests were used to compare the same measures between mLCN formulations as we had no pre-existing hypothesis regarding the differences in myotube alignment between conditions.

#### Sample Sizes

Unless otherwise included in the relevant methods section, exact values of n are reported here. For the plots in **Figures 1E, 1F, S3, and S4A**: n = 3 (Glass D3, Glass D5, iLCN-0.5 D3, iLCN-0.5 D5, iLCN-0.75 D3, mLCN-0.5 D3, mLCN-0.5 D5, mLCN-0.75 D3, mLCN-0.75 D5, NanoSurface D5), 4 (iLCN-0.75 D5, iLCN-1.0 D3, iLCN-1.0 D5, NanoSurface D3), or 5 (mLCN-1.0 D3, mLCN-1.0 D5) independent samples. For the plots in **Figures 2C, 2D, and S5**, n = 5 non-overlapping fields of view across three independent mLCNs; n = 4 isolated fields of view across two independent iLCNs. For the plots in **Figure 2F**, n = 3 independent mLCNs. For the plots in **Figure 3**, n = 3 independent mLCNs (panel A), n = 2 independent mLCNs (panel B), n = 5 independent mLCNs (panel C), and n = 3 independent mLCNs (panel D). For the plots in **Figure 4**, n = 2 independent mLCNs each for ‘– bleb’ and ‘+ bleb’ conditions. For the plots in **Figure S4C, S4D, S4F, and S4G,** n = 2 independent samples each for mLCN-1.0 and NanoSurface substrates.

