## Supplemental Information for "Stiffness anisotropy coordinates supracellular contractility driving long-range myotube-ECM alignment"

| | mLCN-0.5 | mLCN-0.75 | mLCN-1.0 |
|---|---|---|---|
| C6M (mol fraction acrylate) | 0.5 | 0.75 | 1 |
| Hexylamine (mol fraction amine) | 0.5 | 0.25 | 0 |
| I-651 (wt%) | 2 | 2 | 2 |
| BHT (wt%) | 1 | 1 | 1 |

**Table S1. LCN chemical formulations, related to Figure 1.**

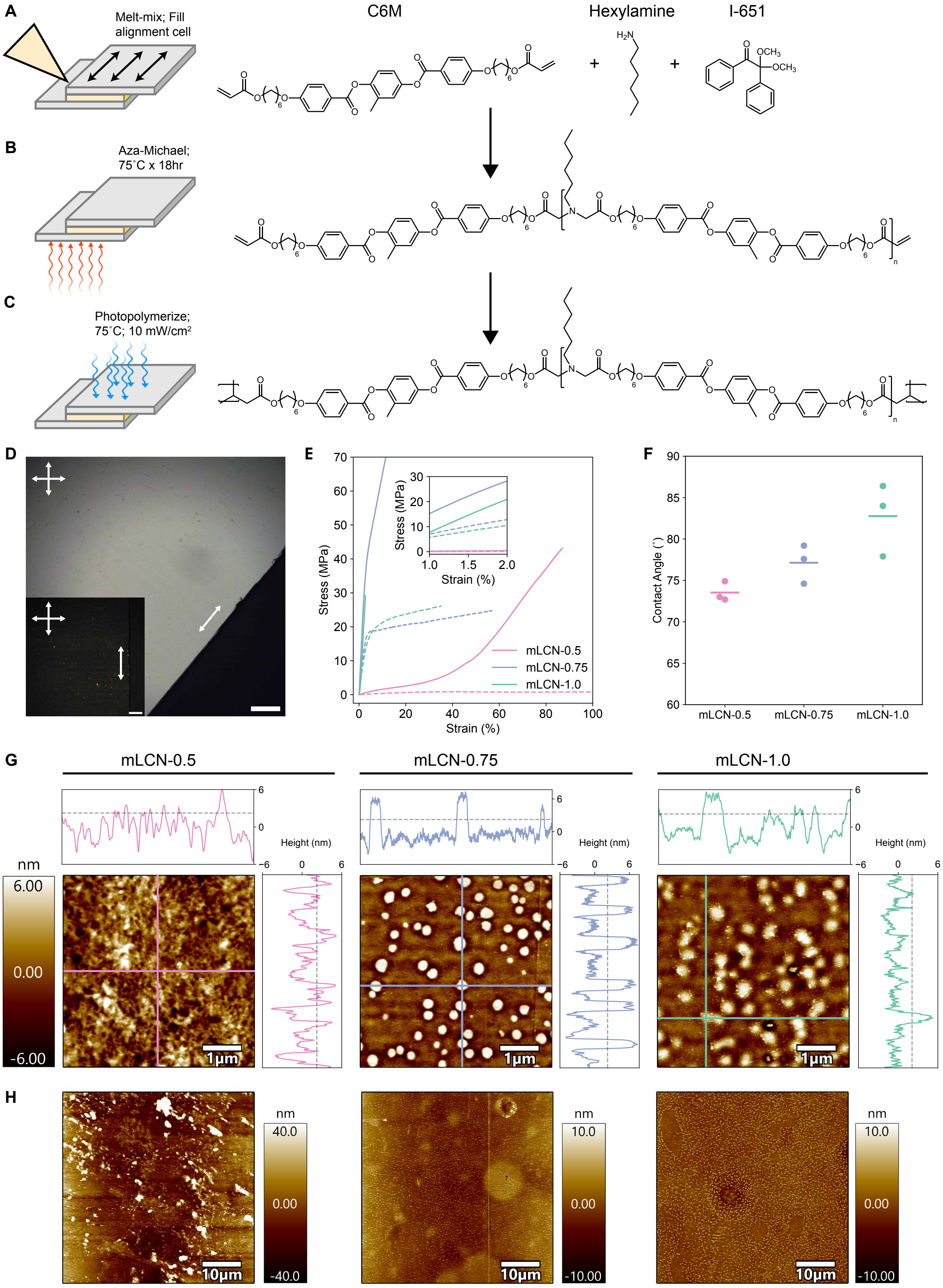

A
Melt-mix; Fill alignment cell
C6M
Hexylamine
I-651
B
Aza-Michael; 75˚C x 18hr
C
Photopolymerize; 75˚C; 10 mW/cm2
D
E
Stress (MPa)
Strain (%)
mLCN-0.5
mLCN-0.75
mLCN-1.0
F
Contact Angle (˚)
mLCN-0.5
mLCN-0.75
mLCN-1.0
G
mLCN-0.5
mLCN-0.75
mLCN-1.0
Height (nm)
nm
6.00
0.00
-6.00
1μm
H
10μm
nm
40.0
0.00
-40.0
10.0
-10.00

**Figure S1**. **LCN film preparation and characterization, related to Figure 1**. **(A)** Liquid crystalline monomer C6M, hexylamine, and photoinitiator I-651 are melted and mixed at 85 ˚C and loaded into a glass surface alignment cell. Arrows indicate rubbing direction which corresponds to the nematic director after polymerization. **(B)** C6M reacts with hexylamine to form oligomers through aza-Michael addition reaction at 75 ˚C for 18 hr inside the cell. The molar ratio of C6M to hexylamine determines the molecular weight of the oligomers. **(C)** Acrylate-terminated oligomers are photopolymerized with UV light (10 $mW/cm^2$) for 15 minutes at 75 ˚C to form a mLCN, or at 135 ˚C to form an iLCN. **(D)** Polarized optical microscopy images demonstrate birefringence and confirm alignment of mLCN-1.0. White arrows indicate nematic director and crossed arrows indicate polarizer directions. Scale bars = 100 µm. **(E)** Representative stress-strain curves from tensile testing of mLCN-0.5, -0.75, and -1.0. Inset shows elastic region of stress-strain curves where elastic modulus values were derived. Solid lines: parallel to nematic director; dashed lines: perpendicular to nematic director. **(F)** Contact angle measurements on mLCN-0.5, mLCN-0.75, and mLCN-1.0. Line indicates mean; all replicates shown. **(G)** High resolution AFM z maps of mLCN-0.5 (left), mLCN-0.75 (middle), and mLCN-1.0 (right) reveal surfaces with root mean square (rms) roughness ~2 nm for all samples (dashed lines in line-scan plots adjacent to images). Small, raised defects on mLCN-0.75 and mLCN-1.0 are quantified by the line-scan plots adjacent to the images and are approximately 10 nm tall and 250 nm in diameter. Defects are randomly distributed with respect to the nematic director (horizontal axis). Scale bars = 1 µm. **(H)** Lower magnification AFM z maps of mLCN-0.5 (left), mLCN-0.75 (middle), and mLCN-1.0 (right) demonstrating the random distribution of small raised defects on mLCN-0.75 and mLCN-1.0 in relation to the nematic director (horizontal axis). Scale bars = 10 µm.

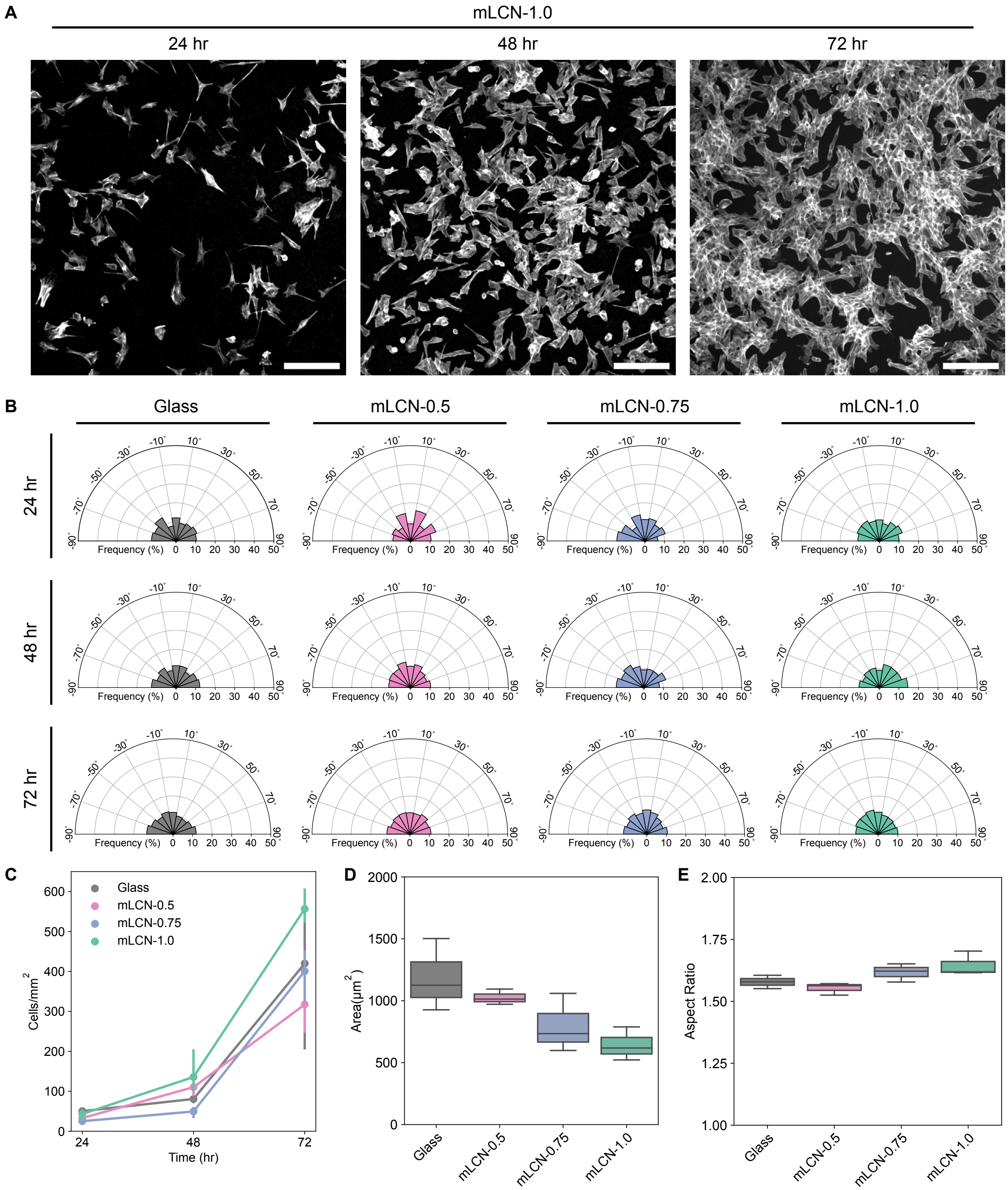


**Figure S2. C2C12 myoblasts have no preference for the nematic director of mLCNs during initial growth phase, related to Figure 1. (A)** Representative images of C2C12 myoblasts stained with phalloidin after 24, 48, and 72 hr of growth on mLCN-1.0. Scale bars = 200 µm. **(B)** Cumulative polar histograms (20˚ bins) of C2C12 myoblast alignment for each substrate after 24, 48, and 72 hr of growth. Data is represented as means of binned frequencies from independent samples. **(C)** Cell density (cells/mm$^2$) for each substrate after 24, 48, and 72 hr of growth. Data is represented as mean ± SD. **(D)** Cell area and **(E)** aspect ratio for each substrate after 72 hr of growth. For the box plots in **(D)** and **(E)**, the box limits extend from the 25$^{th}$ to 75$^{th}$ percentiles; the horizontal line indicates the median value; the whiskers extend by 1.5x the inter-quartile range.

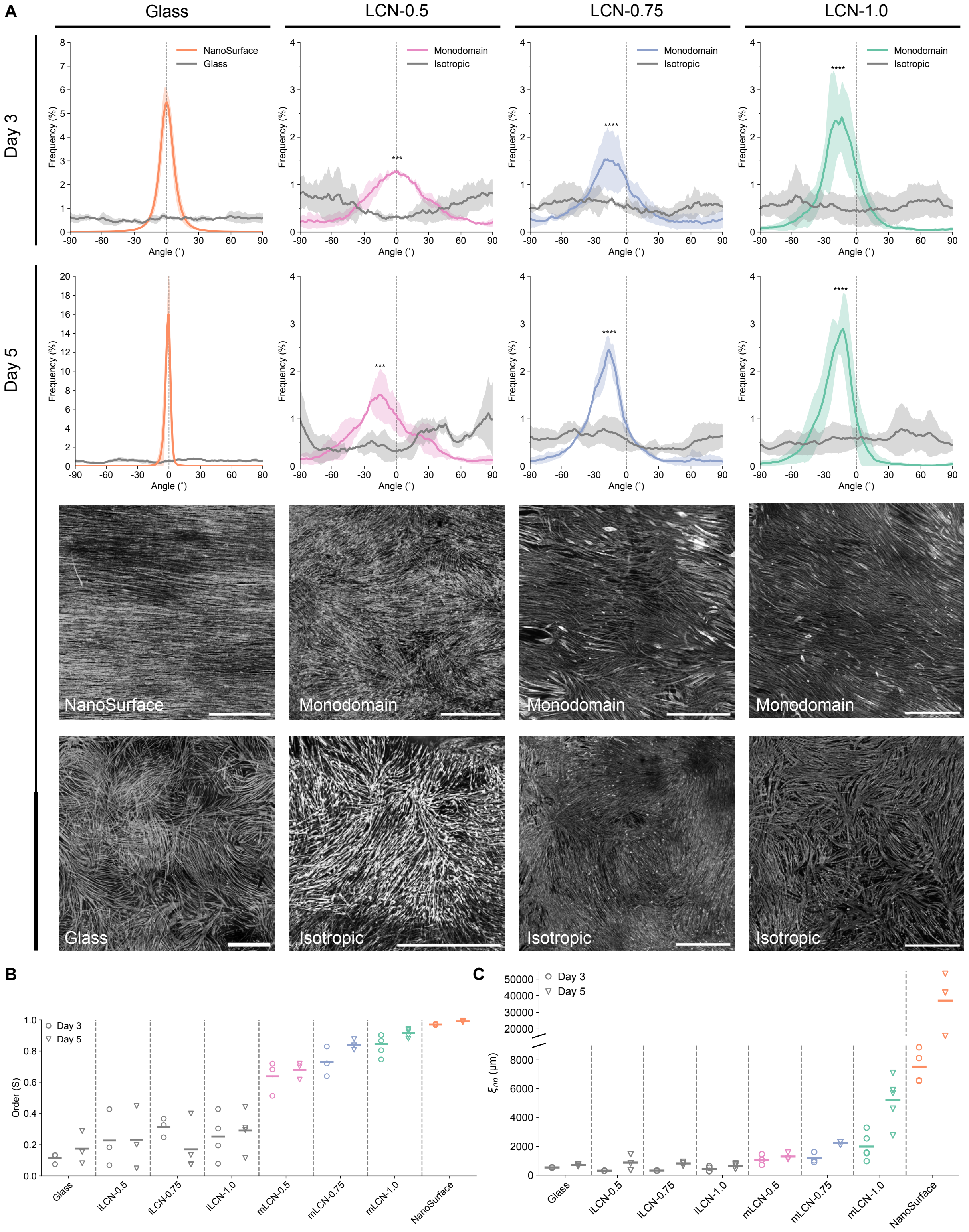
A
Glass
LCN-0.5
LCN-0.75
LCN-1.0
Day 3
Day 5
NanoSurface
Glass
Monodomain
Isotropic
Frequency (%)
Angle (˚)
***
****
NanoSurface
Monodomain
Monodomain
Monodomain
Glass
Isotropic
Isotropic
Isotropic
B
Order (S)
Day 3
Day 5
Glass
iLCN-0.5
iLCN-0.75
iLCN-1.0
mLCN-0.5
mLCN-0.75
mLCN-1.0
NanoSurface
C
$\xi_{nn}$ (μm)
Day 3
Day 5

**Figure S3. Myotube alignment is influenced by the degree of mLCN stiffness anisotropy, related to Figure 1.** **(A)** Frequency distribution plots of myotube alignment on isotropic (gray) and monodomain (colored) LCNs, and glass (gray) or NanoSurface (orange) after 3 (row 1) and 5 (row 2) days of differentiation. Data are represented as mean ± SD. Row 3, 4: Representative images of myosin II heavy chain staining after 5 days of differentiation on NanoSurface, gelatin-coated glass, monodomain LCNs and isotropic LCNs. Scale bars = 1000 µm. Images of LCN-1.0 (monodomain and isotropic) are duplicated from **Figure 1D** for completeness. **(B)** Myotube orientation-order parameter (S) and **(C)** nematic correlation length (µm) on all substrates after 3 and 5 days of differentiation. Line indicates mean; all replicate values shown.

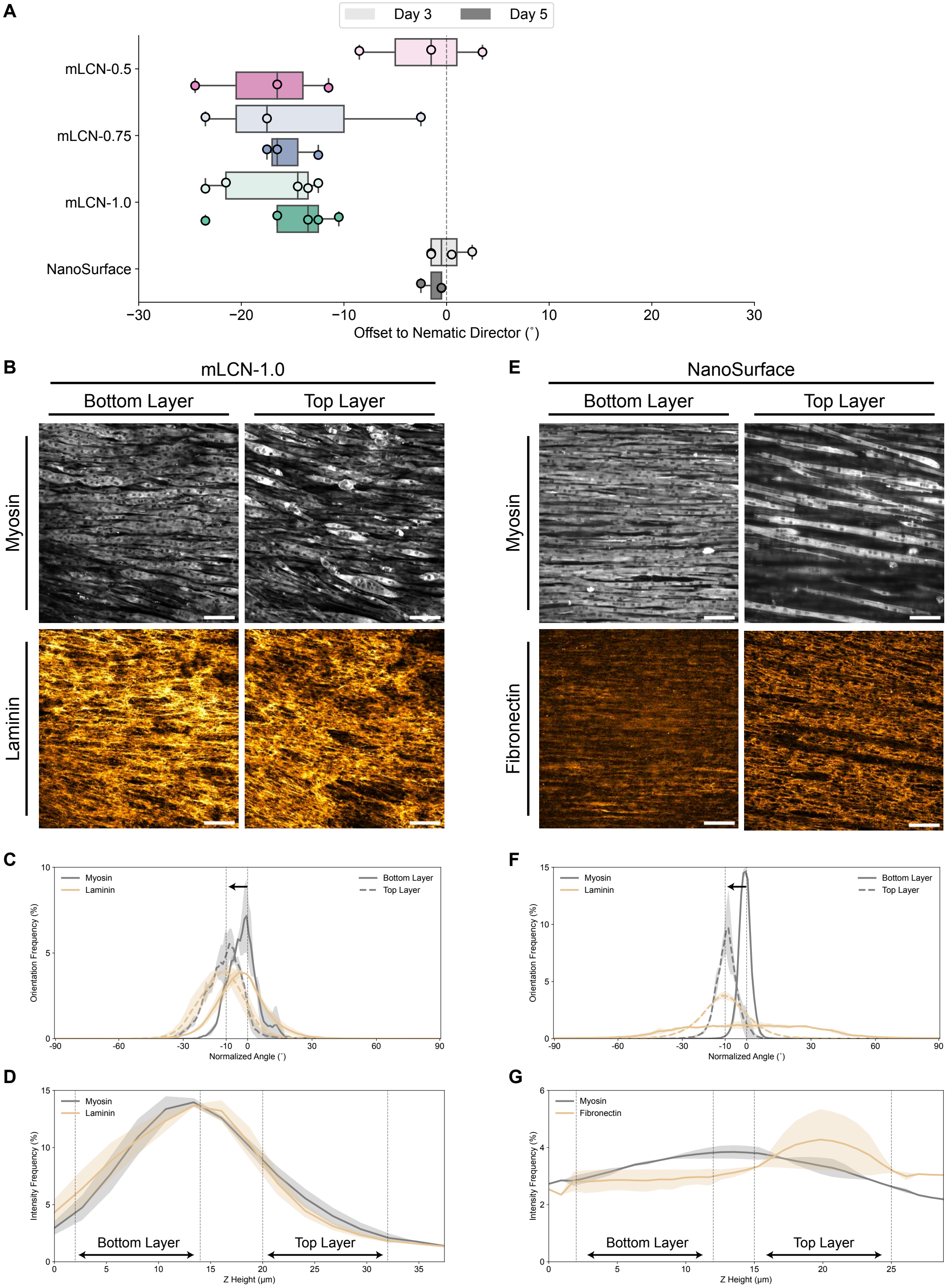

A
Day 3
Day 5
mLCN-0.5
mLCN-0.75
mLCN-1.0
NanoSurface
−30
−20
−10
0
10
20
30
Offset to Nematic Director (°)
B
mLCN-1.0
Bottom Layer
Top Layer
Myosin
Laminin
E
NanoSurface
Bottom Layer
Top Layer
Myosin
Fibronectin
C
Myosin
Laminin
Bottom Layer
Top Layer
Orientation Frequency (%)
Normalized Angle (°)
F
Myosin
Laminin
Bottom Layer
Top Layer
Orientation Frequency (%)
Normalized Angle (°)
D
Myosin
Laminin
Intensity Frequency (%)
Bottom Layer
Top Layer
Z Height (μm)
G
Myosin
Fibronectin
Intensity Frequency (%)
Bottom Layer
Top Layer
Z Height (μm)

**Figure S4. Aligned myotubes develop layers with chiral offset to the layer below, related to Figure 1. (A)** Myotube peak alignment offset to the nematic director of mLCNs and groove direction of NanoSurface substrates after 3 and 5 days of differentiation. The box limits extend from the 25$^{th}$ to 75$^{th}$ percentiles; the horizontal line indicates the median value; the whiskers extend by 1.5x the inter-quartile range; all replicates including outliers shown. **(B)** Representative images of myosin II heavy chain and laminin alignment on mLCN in the bottom layer (left) and top layer (right) after 5 days of differentiation. Scale bars = 100 µm. **(C)** Corresponding orientation frequency distribution plot of myosin and laminin alignment in each layer. Arrow indicates offset from bottom layer to top layer. Data is represented as mean ± SD. **(D)** Corresponding intensity frequency distribution plot of myosin and laminin as a function of height from the bottom of the substrate. Data is represented as mean ± SD. **(E)** Representative images of myosin II heavy chain and fibronectin alignment on NanoSurface substrate in the bottom layer (left) and top layer (right) after 5 days of differentiation. Scale bars = 100 µm. **(F)** Corresponding orientation frequency distribution plot of myosin and laminin alignment in each layer. Arrow indicates offset from bottom layer to top layer. Data is represented as mean ± SD. **(G)** Corresponding intensity frequency distribution plot of myosin and laminin as a function of height from the bottom of the substrate. Data is represented as mean ± SD.

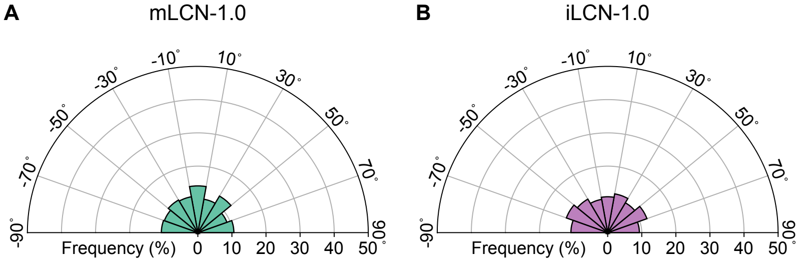


**Figure S5. C2C12 myoblast axis of division is isotropic during proliferative phase, related to Figure 2.** Cumulative polar histograms (20˚ bins) of C2C12 myoblast division angle throughout the proliferative phase on **(A)** mLCNs and **(B)** iLCNs. Data are represented as means of binned frequencies from independent samples.

### Supplemental Video Legends

**Video S1. Collective myoblast polarization and cellular flows on mLCNs and iLCNs, related to Figure 2.** Top: Timeseries of C2C12s on mLCN (left) and iLCN (right) stained with SiR-Actin from $t_a$ = 12–72 hr. Bottom: PIV timeseries corresponding to timeseries above. Vectors are autoscaled within each frame and overlaid on a heatmap of velocity magnitude ranging from 0–5 μm/hr. Frames that are dimmed were excluded from PIV analysis due to stage motion artifacts.

**Video S2. Collective cellular flows annihilate topological defects driving global myotube ordering on mLCNs, related to Figure 2.** Top left: Timeseries of C2C12s on mLCN stained with SiR-Actin from $t_a$ = 48–120 hr. Bottom left: PIV timeseries where vectors are autoscaled within each frame and overlaid on a heatmap of velocity magnitude ranging from 0–8 μm/hr. Top right: Timeseries of orientation vector field line integral convolution demonstrating annihilation of defects during collective cellular flows. Bottom right: Timeseries of local orientation-order parameter heatmap ranging from S = 0–1 demonstrating local regions of disorder (black) that become ordered (white) during collective cellular flows.

**Video S3. Inhibition of cellular contractility with blebbistatin abrogates collective cellular flows and prevents global myotube ordering, related to Figure 4.** Top: Timeseries of C2C12s on mLCN stained with SiR-Actin without (left) or with (right) addition of blebbistatin from $t_a$ = 0–84 μm/hr. Images were taken every 3 hours. Blebbistatin is first added at $t_a$ = 12 hr and media is replaced every 24 hours after. Bottom: PIV timeseries corresponding to timeseries above. Vectors are autoscaled within each frame and overlaid on a heatmap of velocity magnitude ranging from 0–7 μm/hr. Frames that are dimmed were excluded from PIV analysis due to stage motion artifacts.

**Supplemental Note 1**

When analyzing maximum intensity projections of myotubes grown on mLCNs, we observed that the dominant orientation was consistently offset in a clockwise (CW) direction from the nematic director **(Figure S3A)**. The magnitude of this offset varied between ~10–20° after five days of differentiation on all mLCNs **(Figure S4A)**. Prior reports have shown that C2C12 myoblasts demonstrate chiral orientation on fibronectin micropatterns due to chiral actin reorganization and cellular jamming.[S1,S2] When cultured in thin, rectangular fibronectin micropatterns, C2C12 myoblasts were found to exhibit a similar chiral offset.[S3] Uniquely, when we imaged myotube cultures on mLCNs after five days of differentiation with greater axial resolution, we identified distinct myotube layers with a chiral offset to the underlying layer **(Figure S4B)**. The bottom layer, closest to the substrate surface, was typically aligned with the nematic director. The top layer was ~10° CW offset from the layer below **(Figure S4C)**. We then considered whether this chiral layering was unique to mLCNs or could be replicated on the grooved NanoSurface substrate. Indeed, we found that a second layer of myotubes emerged on top of the layer in contact with the nanogrooves after five days of differentiation and exhibited a similar ~10° CW offset **(Figures S4E and S4F)**. Thus, regardless of the mechanism of alignment (stiffness anisotropy vs. contact guidance), the inherent chirality of C2C12 myoblasts is retained upon fusion into myotubes and extends into the third dimension. This chiral myotube layering also explains why we observed a CW shift in the alignment frequency when analyzing the orientation of maximum intensity projections in **Figure S3**.

Given the importance of reciprocal cell-ECM dynamics in driving global myotube alignment on mLCNs **(Figures 3 and 4)**, we also assessed the ECM architecture with confocal microscopy. We found that laminin staining on mLCNs matched both the chiral twist and density of signal (i.e., fluorescence intensity) of the myotube layers (**Figures S4B–D**). Fibronectin alignment on the NanoSurface substrate also matched the chiral twist of the myotubes. However, fibronectin was surprisingly sparse in the bottom layer of myotubes in contact with the grooves. Fibronectin density only increased in the top layer of myotubes above the nanogrooves **(Figures S4E–G)**. This suggests that nascent ECM deposition and remodeling is required for the evolution of chiral myotube layers and maintenance of long-range alignment once myotubes escape contact with topographic features that enforce alignment. These findings also indicate that contact guidance-mediated myotube alignment occurs independent of reciprocal cell-ECM interactions, again highlighting the importance of reciprocal cell-ECM dynamics in maintaining long-range myotube alignment and chiral layering on mLCNs that lack topography.

While the evolution of chiral layers was not the focus of this study, we can still comment on some possible explanations and mechanisms. The chiral offset of C2C12s cultured on thin, rectangular fibronectin micropatterns was found to be a result of the emergence of a shear stress field due to appositional migration on opposing edges of the micropattern.[S3] However, this effect decayed with increasing width of the micropattern and did not result in global order for widths greater than 1000 μm, limiting translation of these results to our studies on mLCNs which were at minimum 3x3 mm. The most relevant study to date found that C2C12 myoblasts evolve into a multilayered culture at stationary +1/2 topological defects.[S4] In that report, the top layer develops a 90˚ offset to the bottom layer. Given the presence of multiple topological defects in our cultures before collective cellular flows emerge, it is feasible that these myoblasts initially develop the same 90˚ offset but are then reoriented to a final offset of ~10–20˚ during the collective cellular flows that align myotubes with the stiffest direction. In another study, C2C12 myoblasts cultured on circular fibronectin micropatterns of the same size scale as the nematic correlation length of C2C12 myoblasts develop into 3D cellular mounds.[S5] These mounds were formed by a combination of active anisotropic stresses and collective cellular flows, with myoblasts exhibiting an increasing chiral offset to the micropattern boundary with each additional layer. Regardless of the mechanism of chiral layering in our study, it is certainly a result of the interplay between the active anisotropic stresses that govern collective cellular dynamics and dynamic cell-ECM interactions that maintain 3D multicellular morphology.

S4. Sarkar T, Yashunsky V, Brézin L, et al. Crisscross multilayering of cell sheets. *PNAS*

*Nexus*. 2023;2(3):1-11. doi:10.1093/pnasnexus/pgad034

S5. Guillamat P, Blanch-Mercader C, Pernollet G, Kruse K, Roux A. Integer topological defects organize stresses driving tissue morphogenesis. *Nat Mater*. 2022.
